# Reviewing the consequences of genetic purging on the success of rescue programs

**DOI:** 10.1101/2021.07.15.452459

**Authors:** Noelia Pérez-Pereira, Armando Caballero, Aurora García-Dorado

## Abstract

Genetic rescue is increasingly considered a promising and underused conservation strategy to reduce inbreeding depression and restore genetic diversity in endangered populations, but the empirical evidence supporting its application is limited to a few generations. Here we discuss on the light of theory the role of inbreeding depression arising from partially recessive deleterious mutations and of genetic purging as main determinants of the medium to long-term success of rescue programs. This role depends on two main predictions: (1) The inbreeding load hidden in populations with a long stable demography increases with the effective population size; and (2) After a population shrinks, purging tends to remove its (partially) recessive deleterious alleles, a process that is slower but more efficient for large populations than for small ones. We also carry out computer simulations to investigate the impact of genetic purging on the medium to long term success of genetic rescue programs. For some scenarios, it is found that hybrid vigor followed by purging will lead to sustained successful rescue. However, there may be specific situations where the recipient population is so small that it cannot purge the inbreeding load introduced by migrants, which would lead to increased fitness inbreeding depression and extinction risk in the medium to long term. In such cases, the risk is expected to be higher if migrants came from a large non-purged population with high inbreeding load, particularly after the accumulation of the stochastic effects ascribed to repeated occasional migration events. Therefore, under the specific deleterious recessive mutation model considered, we conclude that additional caution should be taken in rescue programs. Unless the endangered population harbors some distinctive genetic singularity whose conservation is a main concern, restoration by continuous stable gene flow should be considered, whenever feasible, as it reduces the extinction risk compared to repeated occasional migration and can also allow recolonization events.

---

Genetic rescue is the reduction of the extinction probability of endangered populations through the introduction of migrant individuals. Genetic rescue programs have been successful in multiple occasions (Vilà et al. 2003; Fredrickson et al. 2007; Johnson et al. 2010; Åkesson et al. 2016; Weeks et al. 2017; Hasselgren et al. 2018; Ralls et al. 2020), and are considered a promising strategy in Conservation Biology (Waller 2015; Tallmon 2017). However, most information on their consequences refer to a few generations (usually one or two, rarely six, Whiteley et al. 2015). Furthermore, concern has been raised by the extinction of the Isle Royale wolves population, where the genetic contribution of a single migrant wolf from the large mainland population quickly spread in the resident population thanks to the breeding vigor of its offspring, possibly causing an increase in inbreeding and an associated fitness decline that triggered population extirpation (Hedrick et al. 2014, 2017, 2019). Therefore, despite the multiple studies supporting the practice of genetic rescue (Frankham 2015; Kolodny et al. 2019), its consequences in the medium to long term remain uncertain (Hedrick and Fredrickson 2010; Hedrick and García-Dorado 2016; Bell et al. 2019; Kyriazis et al. 2020; Ralls et al. 2020). Here we review the main theoretical aspects behind the impact of inbreeding depression and purging on the long-term success of rescue programs and carry out computer simulations to evaluate the predicted outcome of these programs under some specific scenarios.

## Some background on purging and on its role during genetic rescue

From the genetic point of view, the main determinant of early future extinction of small endangered populations is the inbreeding depression of fitness (O’Grady et al. 2006; Allendorf et al. 2013; Frankham et al. 2014). This is due to the expression, as inbreeding accumulates, of the initial inbreeding load *B*, often interpreted in terms of lethal equivalents (Morton et al., 1956). Here we deal with the inbreeding load *B* ascribed to the recessive deleterious component of many rare detrimental alleles that remains hidden in the heterozygous condition in a non-inbred population (see, e.g., Caballero 2020, Chap. 8), and we do not consider the possible inbreeding load ascribed to overdominance. According to theory, in stable populations the inbreeding load is expected to be larger for populations with larger effective size *N* (see Eq. 13 in García-Dorado 2007), the increase being much more dramatic for more recessive deleterious alleles (García-Dorado 2003; Hedrick and García-Dorado 2016). Thus, a historically large population can be genetically healthy in the sense of showing a high average fitness but, still, its individuals are expected to be heterozygous for many rare (partially) recessive deleterious alleles.

Due to the reduction of *N* in an endangered population, both drift (the dispersion of gene frequencies due to random sampling of alleles) and inbreeding (the increase in homozygosis in the offspring of related individuals) increase through generations. Inbreeding increases the expression of recessive deleterious effects in homozygosis, which produces inbreeding depression but also triggers an increase of selection against the deleterious alleles known as genetic purging. The process is described by the Inbreeding-Purging model (IP model in García-Dorado 2012), according to which the fitness expected by generation *t* (*W_t_*) after a reduction of *N* is predicted as in the classical Morton’s et al. (1956) model, but replacing Wright’s inbreeding coefficient *F* with a purged inbreeding coefficient *g* that is weighed by the ratio *q_t_ /q_0_*, where *q_0_* is the frequency of the deleterious allele in the original non-inbred population and *q_t_* is the corresponding value expected from purging by generation *t*:

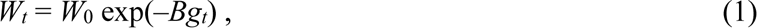

where *W*_0_ and *B* are, respectively, the expected fitness and the inbreeding load in the initial non-inbred population, and where *g_t_* can be predicted as a function of the effective population size *N* and of the recessive component of the deleterious effects, i.e., the purging coefficient *d* which, for a given homozygous effect *s* and dominance coefficient *h*, amounts *d* = *s*(1 – 2*h*)/2. Thus, *B* is the sum of 2*d*(1 – *q*_0_)*q*_0_ over all the sites with segregating deleterious alleles. Similarly, the corresponding inbreeding load at generation *t* (*B_t_*) can be predicted as

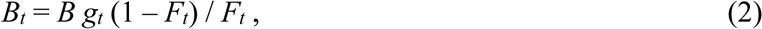

which accounts for the joint reduction of the inbreeding load ascribed to drift and purging, that is faster than under drift alone.

The efficiency of purging can be defined as the proportional reduction of the original deleterious allele frequencies that it is expected to cause, i.e., the expected (*q_0_ -q_t_* )*/q_0_*. Therefore, since the asymptotic value of *g*,

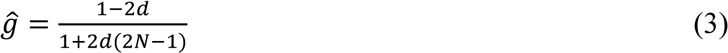

predicts the asymptotic value of *q_t_* /*q_0_* to a good approximation for *Nd* ≥ 1, we predict the efficiency of purging as (1 − *g̃*). This expression accounts for the opposing effects of purging and genetic drift after long-term inbreeding, when all the deleterious alleles responsible for the initial *B* are expected to be fixed or lost. It shows that the efficiency of purging increases with increasing *Nd* being, for any given *d* value, higher in large populations (i.e., under slower inbreeding) than in small ones. As *d* approaches 0, *ĝ* approaches 1, and the role of purging preventing deleterious fixation becomes negligible compared to that of drift. As *Nd* increases, *ĝ* goes to zero, drift becomes irrelevant and the deleterious alleles responsible for *B* in the original non-inbred population are expected to be virtually removed by purging. Figure 1 illustrates that purging occurs faster under faster inbreeding (i.e., for smaller *N*) but it is also less efficient.

**Figure 1.**
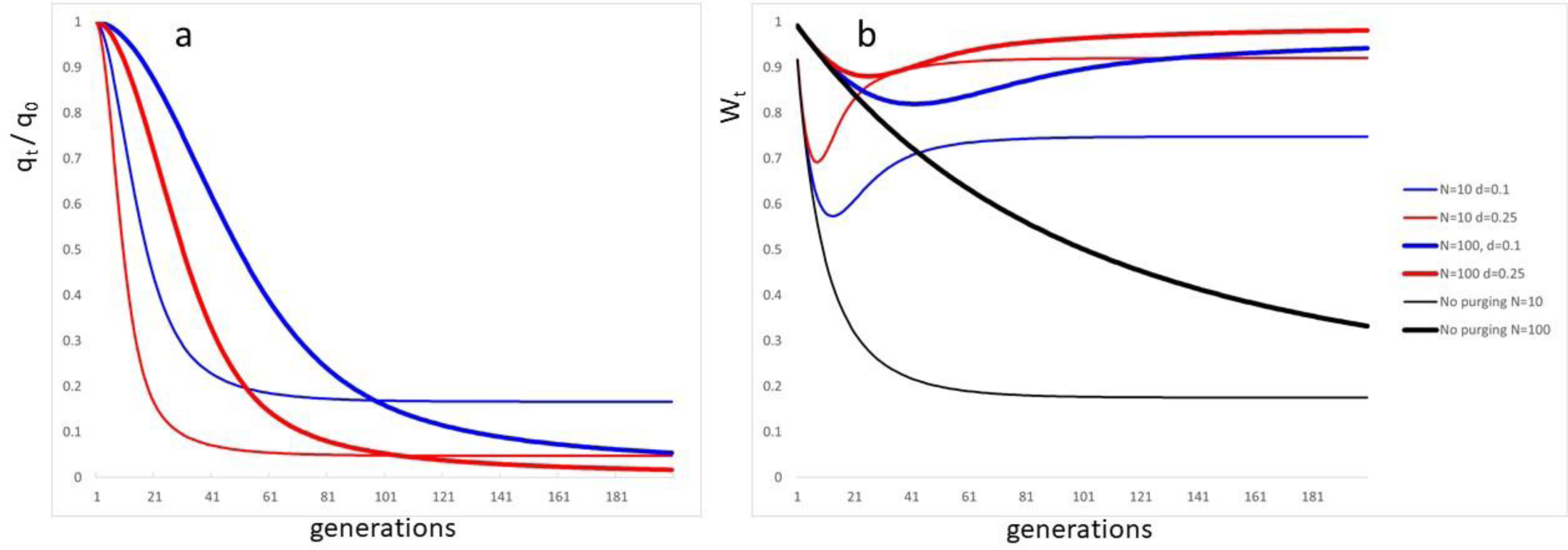
Consequences of purging over 200 generations after the effective population size of an ancestral large population with *W0* = 1 and *B* = 2 drops to *N* = 10 (thin lines) or *N* = 100 (thick lines); blue: *s* = 0.2, *h* = 0, *d* = 0.1; red: *s* = 0.5, *h* = 0, *d* = 0.25; black: neutral (no purging) predictions. a) Average of the frequency of the deleterious alleles of the ancestral population through generations relative to the corresponding initial frequency (*qt*/*q*0, inferred as *gt*/*Ft*). In the absence of selection this average relative frequency would remain equal to 1. However, it is substantially reduced due to purging. The reduction occurs faster for smaller *N*. After some time, an equilibrium is reached where the average relative frequency represents the fraction of ancestral deleterious allele that become fixed because they have not been purged. This asymptotic average frequency is larger for smaller populations, indicating less efficient purging. Purging is quicker and more efficient for larger *d* values; b) Expected average fitness through generations showing initial inbreeding depression and later substantial recovery due to purging, although never up to its ancestral value. A more comprehensive model including non-purging selection and new mutation (the Full Model) can also be found in García-Dorado (2012).

In agreement with the above predictions, there is evidence that purging is able to reduce an important fraction of the inbreeding depression in populations with effective sizes about ten or above, while faster inbreeding (as continued full-sib mating or effective sizes below 10) seems to promote purging just against lethal or severely deleterious mutations (Templeton and Read 1984; Hedrick 1994; Wang et al. 1999; Ávila et al. 2010; Pekkala et al. 2012; Bersabé and García-Dorado 2013; López-Cortegano et al. 2016; Caballero et al. 2017).

Therefore, the success of genetic rescue programs to reduce the extinction risk ascribed to inbreeding depression depends on the balance between two different effects of gene flow. On the one hand, migrants reduce inbreeding, thus causing an increase of fitness that corresponds to reversed inbreeding depression, which is known as hybrid vigor or heterosis (Falconer and Mackay 1996, p. 253; Caballero 2020, p. 196) and is due to the introduction of the beneficial allele at some of the sites where the individuals of the endangered populations were homozygous for the deleterious allele. Obviously, the more purging occurred before migration, the smaller is the inbreeding depression accumulated in the endangered population and the corresponding hybrid vigor induced by the rescue program. On the other hand, migrants bear their own inbreeding load due to partially recessive deleterious alleles hidden in heterozygosis. This hidden inbreeding load may fuel future inbreeding depression in the endangered population, which can be mitigated by purging. Therefore, the success of genetic rescue programs can critically depend on the purging occurring on both the donor population and the endangered recipient one. The impact of a rescue program on the extinction risk also depends on many other factors besides inbreeding and purging, such as the possible advantage due to migrants contributing new adaptive mutations accumulated in the donor population after the isolation of the endangered one, the possible adaptive disruption if the two populations are adapted to different environments, the increased resilience due to restoration of adaptive potential, the introduction of demographic and environmental stochasticity or other factors related to management as the risk of spread of infectious diseases, etc. (Ralls et al. 2020). However, in this review we will focus on genetic purging considering theoretical predictions and available evidence, including new simulation results, to understand its role as a determinant of the success of genetic rescue programs in reducing both inbreeding depression and extinction risk through generations.

## Genetic purging in the donor population

Drawing migrant individuals from large, genetically healthy populations has usually prompted population recovery during a few generations (Whiteley et al. 2015; Ralls et al. 2020). Nevertheless, as explained in the previous section, migrant individuals sampled from a historically large donor population are expected to be heterozygous for many rare (partially) recessive deleterious alleles. Each of these alleles cause slight or no damage on the fitness of migrants and of the offspring they produce when mating individuals of the recipient population. However, they may contribute inbreeding load to the recipient population that can cause an increase of the inbreeding depression in the future. Thus, using large donor populations to rescue very small endangered ones could in theory enhance the risk of extinction from future inbreeding depression.

Therefore, in some cases, migrants from slowly inbred efficiently purged populations (i.e. where inbreeding has accumulated due to effective population sizes above several tens), could be a better alternative in the medium to long term. These migrants can produce hybrid vigor without a substantial increase of the inbreeding load and of the long-term extinction risk, even if leading to smaller gains in genetic diversity and, therefore, in adaptive potential. It has been stated that reliable evidence is required about the superiority of migrants sampled from small populations (Ralls et al. 2020) but, in fact, except when considering just a few generations, reliable evidence is required for the success of rescue using both small and large donor populations.

It needs to be remembered that purging becomes less efficient for smaller populations. Therefore, using migrants from a population that underwent drastic bottlenecking can introduce high genetic load as well as little genetic diversity and adaptive potential, bringing together the worst of both worlds. This seems to have been the case with one of the donor populations used to rescue the endangered Pacific pocket mouse (Wilder et al. 2020). Fortunately, analysis of genomic data can provide inferences on the demographic history (Santiago et al. 2020, and references therein) that may allow the election of a donor population with a record of moderate size allowing for efficient purging.

It has been proposed that the increase of extinction risk ascribed to the deleterious alleles introduced during genetic rescue can be controlled by prioritizing a lower putative load inferred from genomic analysis over a high genetic diversity (Kyriazis et al. 2020; Teixeira and Huber 2021). However, relying on the ability to identify the mutations that are responsible for a main fraction of the fitness load is not free of perils (Kardos and Shafer 2018; Ralls et al. 2020; García-Dorado and Caballero 2021). For example, the number of putatively deleterious variants per genome has not been found to be a reliable predictor of inbreeding depression in island populations of foxes and wolves (Robinson et al. 2018). Similarly, the Homozygous Mutation Load, defined by Keller et al. (2011) as the number of homozygous loci for rare alleles carried by an individual, used as a proxy of the fitness load due to homozygous (partially) recessive deleterious mutations, has only a moderate expected correlation with phenotypic values of individuals in large populations (Caballero et al. 2020). The recent evidences that purging can reduce different genomic proxies for the fitness genetic load (Xue et al. 2015; Robinson et al. 2018; Van der Valk et al. 2019; Grossen et al. 2020) suggest that genomic analyses could be helpful to infer the inbreeding load at the population level, but this is more likely to be useful in identifying suitable donor populations than optimal migrant individuals. However, even identifying the best donor population is not straightforward based on genomic information, as between population differences in fitness inbreeding load can be mainly due to a very small fraction of the annotated alleles of any deleterious category. For example, considering two populations with a common origin but different demographic history, one of them could have purged many more mildly deleterious alleles than the other during a long evolutionary period, but the other one could have purged a few more severely deleterious alleles during a recent shorter period with a relatively smaller size. Thus, the population with the smallest count of putatively deleterious alleles per genome is not necessarily the population with the lower fitness or the smaller fitness inbreeding load.

Thus, when the donor’s inbreeding load is a concern (see next section), it can be safer preferring donor populations that have gone through a period of moderate effective size allowing for purging in the past than choosing migrant individuals on the basis of their low burden of putatively deleterious alleles. Adaptive potential could be further improved by using different donor populations if available. For example, some of the populations of Canadian lynx in eastern North America could need to be rescued in the future due to global warming preventing natural migration through natural ice bridges. Then, migrants could be sampled from the several peripheral populations in eastern Canada that are under continuous partial isolation instead of from the large mainland Canada population (Koen et al. 2015).

A relevant question is in which situations the load introduced by non-purged migrants can be more harmful than the inbreeding depression they remove. The answer depends on the purging processes that take place in the recipient endangered population, analyzed in the next section.

## Purging occurring in the recipient population

### Theoretical arguments

Let us now think of a rescue program where the donor population has a long history of large effective population size, so that it can be considered genetically healthy (i.e. it shows little reduction of mean fitness from segregating and fixed deleterious alleles) but it is non purged (i.e., it hides large inbreeding load). The success of the rescue program depends on the balance between the inbreeding depression of the endangered population that is intended to be reversed by the migrant gene flow and on the future depression that can arise from the load concealed in the migrant individuals.

In the past, when the endangered population began to shrink, the inbreeding load *B* of the ancestral non-endangered population fueled both inbreeding depression and purging. As predicted by Equations (1) and (2), the slower was the increase of inbreeding during that process (i.e., the larger was *N*) the slower but more efficient was purging (Eq. 3 and Figure 1). This means that, the slower was the process that led the recipient population to its current inbreeding level: a) the smaller is its expected inbreeding depression, so that less hybrid vigor is expected after migration; b) the smaller inbreeding load it hides, so that migrant individuals coming from a large population are more likely to carry more (partially) recessive deleterious alleles than resident ones. In addition, if the recipient population is very small by the time of migration and does not experience a quick demographic recovery, the load introduced by migrant individuals will not be efficiently purged. The result is that, if the recipient population has a history of slow inbreeding previous to migration but remains very small after that, migration events could in some cases reduce fitness in the medium to long term. On the contrary, migration events are expected to be particularly beneficial for populations with a history of drastic bottlenecking that have recovered a moderate size allowing future purging.

### A simulation illustration

To assess the relevance of the purging occurring in the endangered recipient population we performed a simulation analysis that is summarized in the Boxes below and is reported with more detail in the Supplementary Material.

#### BOX 1. Purging and fitness rescue

First, we simulated a large non-threatened population of *N* = 10^4^ individuals for 10^4^ generations to approach the mutation-selection-drift equilibrium. Then, smaller threatened populations were derived and different scenarios were simulated as shown in Fig. Box 1.1. In a first phase, threatened populations with different sizes (*N*_1_ = 4, 10 or 50) were maintained for *t* = *N*_1_ generations (e.g., up to generation 50 for populations with *N*_1_ = 50, etc.), so that the average inbreeding coefficient was *F* ≍ 0.4. Then, in a second phase with population size *N*_2_, each threatened population was maintained with the same constant size (*N*_2_ = *N*_1_), or with a different size, and entered or not a genetic rescue program. Each two-phase scenario is denoted by the corresponding population sizes (*e.g*., 50-10 for *N*_1_ = 50 and *N*_2_ = 10). Rescue consisted of the addition of males randomly sampled from the large *N* = 10^4^ population. Between 500 and 2,500 replicated rounds were simulated. The number of individuals introduced during each migration event was five for lines with *N*_2_ = 50 and one for lines with *N*_2_ = 10 or *N*_2_ = 4. Regarding the number of migration events, we considered four strategies: i) a single event; ii) two events with an interval of five generations; iii) periodic migration every five generations; and iv) the “one migrant per generation” (OMPG) strategy, that is widely recommended to retain connectivity in metapopulation management (Mills and Allendorf 1996). All the sizes considered for the threatened populations (4, 10 and 50) correspond to the IUCN Red List category of Critically Endangered or Endangered according to Criterion D (IUCN 2012).

Non-recurrent deleterious mutations occurred at rate *λ* = 0.2 per gamete and generation in free recombining sites, with fitness 1, 1 – *sh*, 1 – *s* for the wild-type homozygote, the heterozygote and the homozygote for the mutant allele, respectively. The inbreeding load *B* was calculated as the sum of *s*(1 – 2*h*)*pq* for all selective sites, where *q* and *p* = 1 – *q* are the frequencies of the mutant and wild allele, respectively (Morton et al. 1956). The homozygous deleterious effect *s* was sampled from a gamma distribution with mean *s̅* = 0.2 and shape parameter *β* = 0.33, and the dominance coefficient *h* was obtained from a uniform distribution between 0 and e^(-*ks*)^, where *k* is such that the average *h* value is ^ℎ̅^ = 0.283, so that alleles that are more deleterious are expected to be more recessive (Caballero and Keightley 1994). Sampled *s* values larger than 1 were assigned a value *s* = 1 so that the mutational model generates a lethal class. Fitness was multiplicative across loci. The rationale for this mutational model is discussed below. In addition, neutral mutation was simulated to obtain estimates of neutral genetic diversity. One half of the individuals were assigned to each sex, and they were randomly chosen to breed according to their fitness and mated panmictically allowing for polygamy. A more detailed description is given in the Supplementary Material.

**Figure Box 1.1.**
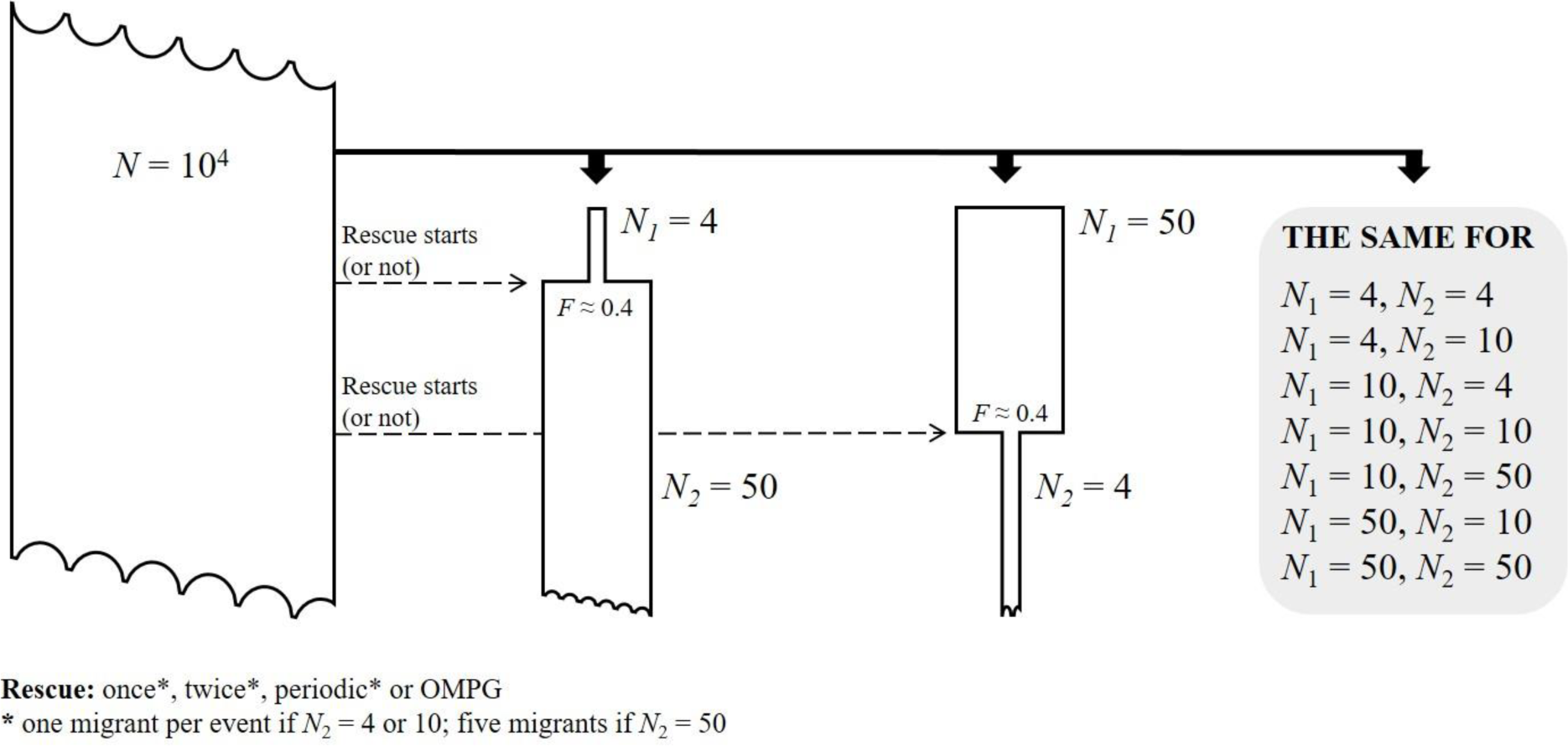
Simulation scheme. A small number of individuals (*N*1) is sampled from the base population to found a threatened population. After *t* = *N*1 generations the population size can either be maintained (*N*2 = *N*1) or changed, and the population can enter, or not, a rescue program. Note that the time progresses downwards and that a waved edge is represented to indicate that the population was maintained with the same size before or after the time represented in the figure.

Fig. Box 1.2 gives population fitness *w* and inbreeding load *B* averaged over replicates in a representative set of scenarios, always computed excluding migrants. Complementary results are given in the Supplementary Material. These results, analyzed in more detail in the main text, illustrate how, after the hybrid vigor occurred in the generations following migration events, some fitness rescue persists over generations when *N*_1_ ≤ *N*_2_ but a fitness disadvantage can be generated compared to “non-rescued” populations when *N*_1_ > *N*_2_. It also shows that periodic migration induces strong fitness oscillations.

**Figure Box 1.2.**
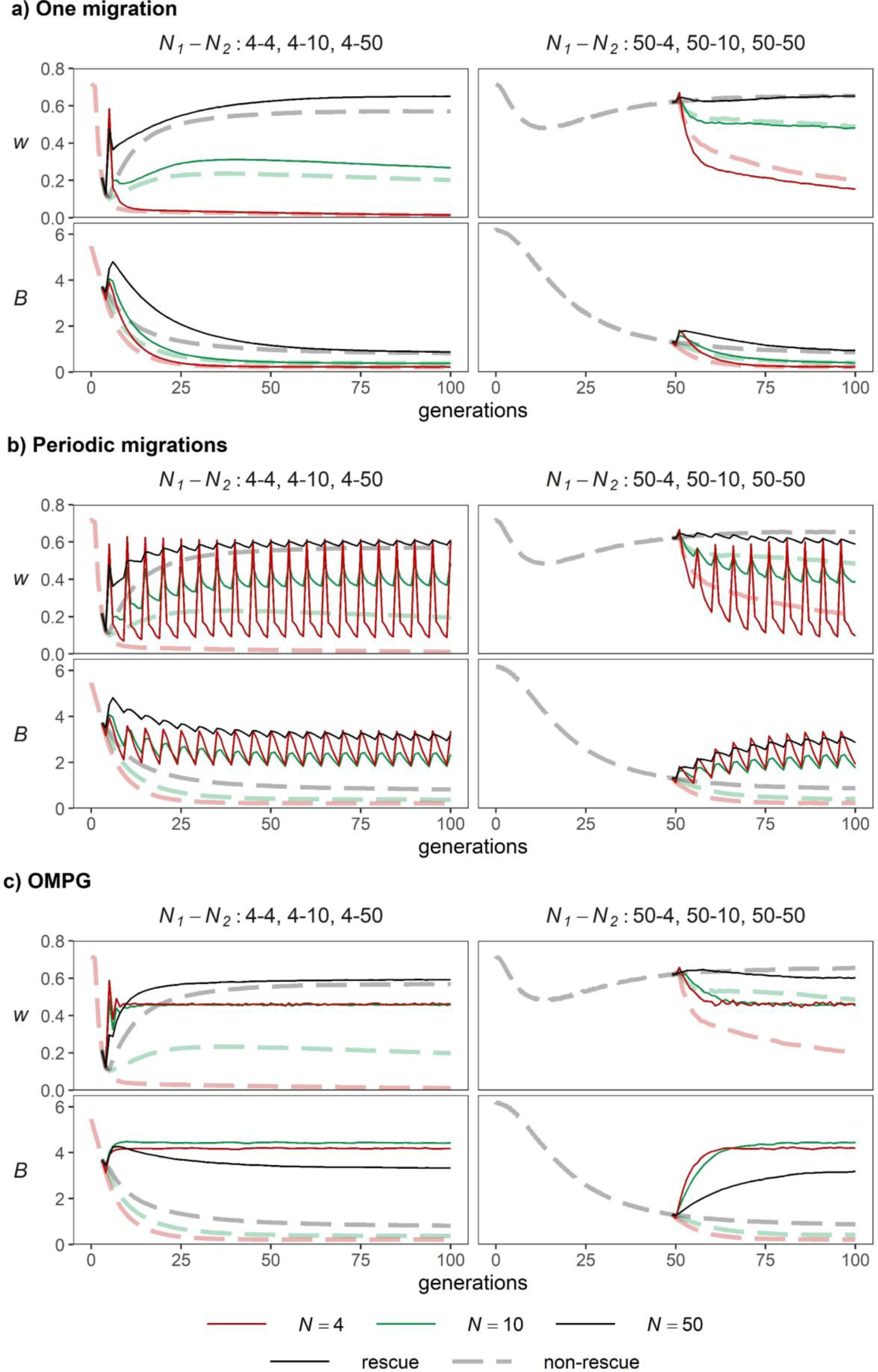
Evolution of average fitness (*w*; upper panels) and inbreeding load (*B*, lower panels) for endangered populations under different demographic and migration scenarios. Demographic scenarios are coded as *N*1-*N*2, *N*1 indicating the population size during phase 1, and *N*2 during phase 2 (red, green and black lines for population sizes 4, 10 or 50, regardless the phase). Light dashed lines represent non-rescue lines. Dark solid lines represent lines entering a rescue program starting at generation *t = N*1. (a) One unique migration of 5 males in lines *N*2 = 50, and of 1 male otherwise. (b) Periodic migrations of 5 males every five generations in lines *N*2 = 50, and of 1 individual otherwise. (c) One migrant male per generation (OMPG).

Figure Box 1.3 illustrates the between-population fitness variability introduced by periodic migration (upper panels) or OMPG (lower panels) for populations with sizes *N*_1_ = 50, *N*_2_ = 4. The panels on the left give the evolution of mean fitness for each of five randomly sampled populations under a non-rescue program; the panels on the right are for populations under rescue. Although in this 50-4 case periodic migration and OMPG increased long-term fitness if averaged over generations (Fig. Box 1.2), both strategies introduce important fitness variability between populations, which adds to the temporal oscillations in the case of periodic migration. This figure illustrates that every input of migrants in a small population can lead to a dangerous inbreeding depression after a few generations.

**Figure Box 1.3.**
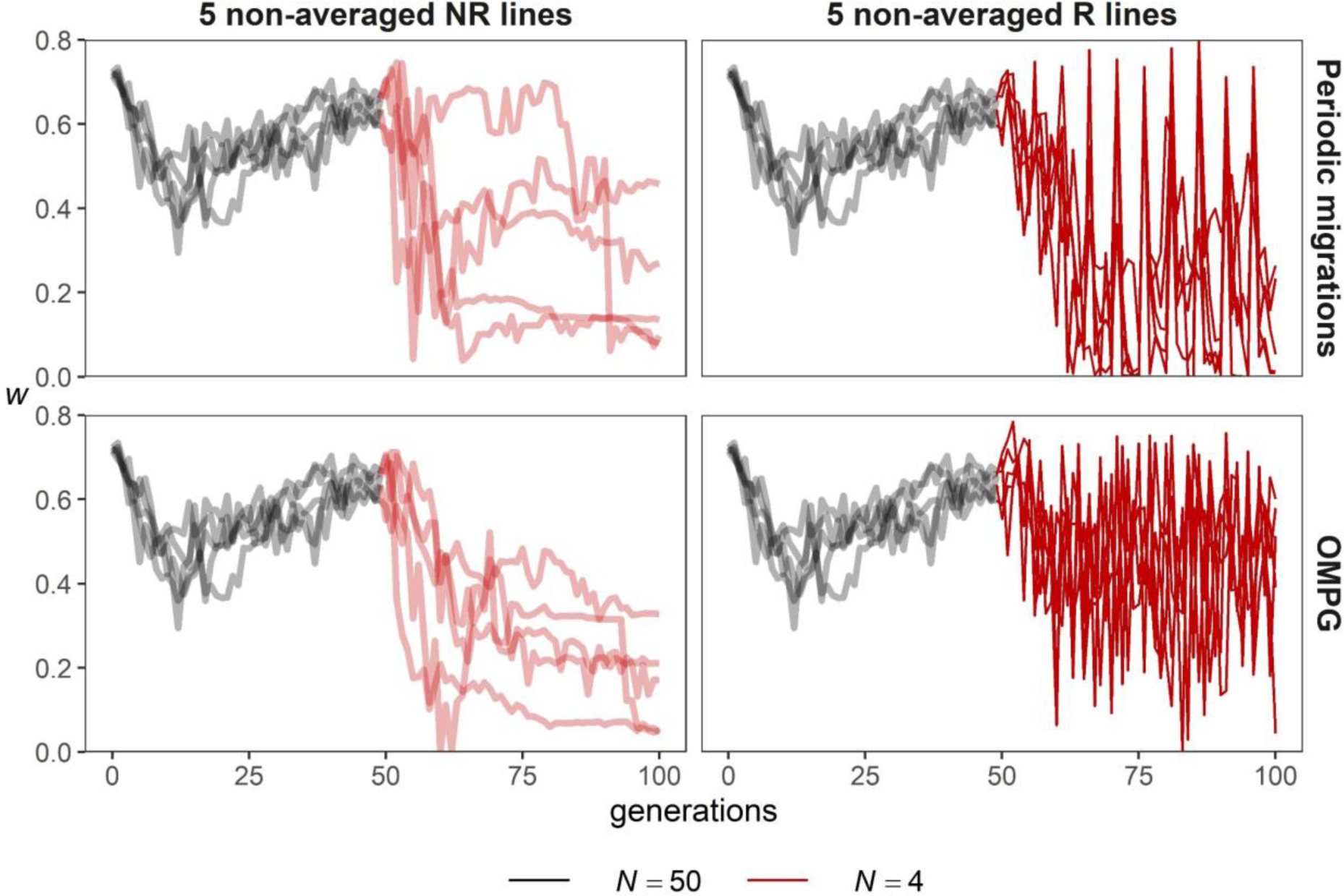
Genetic stochasticity for average fitness (*w*) on sets of five random populations from those described in Fig. Box 1.2 for a scenario with population size *N*1 = 50 during phase 1 and *N*2 = 4 during phase 2 (rescue starting at generation 50). Left panels: populations under no rescue program. Right panels: populations under a rescue program starting at generation 50 (upper panels for periodic migration every 5 generations, lower panels for OMPG).

Results in Fig. Box 1.2 give results averaged over replicates that illustrate the expected evolution of average fitness (*w*) and inbreeding load (*B*) over generations. They show that, under no rescue program, the expected fitness of threatened populations declined in the early generations and partially recovered a few generations later due to genetic purging. The decline was more dramatic and the recovery was poorer or non-existent in the smaller populations, as expected from less efficient purging. The inbreeding load also declined due to both genetic purging and drift. For the larger populations, this decline was much faster than that of the genetic diversity (*H*) for neutral loci, due to efficient purging (Supplementary Material Figs. S1-S4). Finally, a new mutation-selection-drift balance was attained where *B* depended just on *N*_2_, while fitness depended on the size of the population through the whole period considered and continued to decline in the smaller populations due to the continuous fixation of deleterious mutations.

The introduction of migrants into the threatened populations always resulted in the increase of expected fitness (hybrid vigor) in the next generation, at the cost of an increase of the inbreeding load. Under occasional migration (one or two migration event), *B* declined after this initial increase, approaching the same equilibrium values as those of populations under no rescue program. In contrast, under periodic migration and OMPG, *B* oscillated around a plateau for values larger than those achieved under no rescue program.

The hybrid vigor after occasional migration was followed by new inbreeding depression to the point that, for *N*_1_ > *N*_2_, where purging is more efficient before than after genetic flow, the expected fitness soon dropped to values persistently smaller than those of non-rescued lines.

Periodic migration every five generations produced a persistent rescue effect on expected fitness in all the same scenarios as occasional migration, as well as in all the cases with *N*_2_ = 4 including those with *N*_1_ > *N*_2_. However, it led to a strongly oscillatory behavior of the expected fitness (Fig. Box 2.1 and S3).

OMPG also improved expected fitness in all cases with the exception of 50-50 and 50-10, where it induced a slight disadvantage that still persisted by generation 100 (Fig. Box 2.1 and S4). The average advantages were stable through generations and were larger than under periodic migration.

These results are in general agreement with the qualitative predictions presented above (see *Theoretical arguments*). But, still, the main purpose in conservation genetics is not maximizing the expected average fitness but preventing population extinction (Bell et al. 2019). Fig. Box 1.3 shows the evolution of average fitness for five randomly sampled non rescued lines and for five lines under periodic migration or OMPG for the 50-4 scenario. It illustrates that rescue events introduce temporal instability for the fitness of each line and increase the between-lines fitness variance which, particularly under periodic migration, often leads to null or very small fitness values that would imply population extinction. Therefore, a positive effect of rescue on expected fitness does not imply a reduction in extinction risk.

#### BOX 2. Consequences on population extinction

Here we show extinction results corresponding to the scenarios described in Box 1. In these simulations, a population was considered extinct when the average fitness (*w*) of males and/or females was less than 0.05 and/or when there were only breeding males or only breeding females (counted after migration when appropriate). Figure Box 2.1 shows the percentage of initial populations that survived through generations. Genetic parameters averaged over surviving lines are given in Figs. S5-S8.

**Figure Box 2.1.**
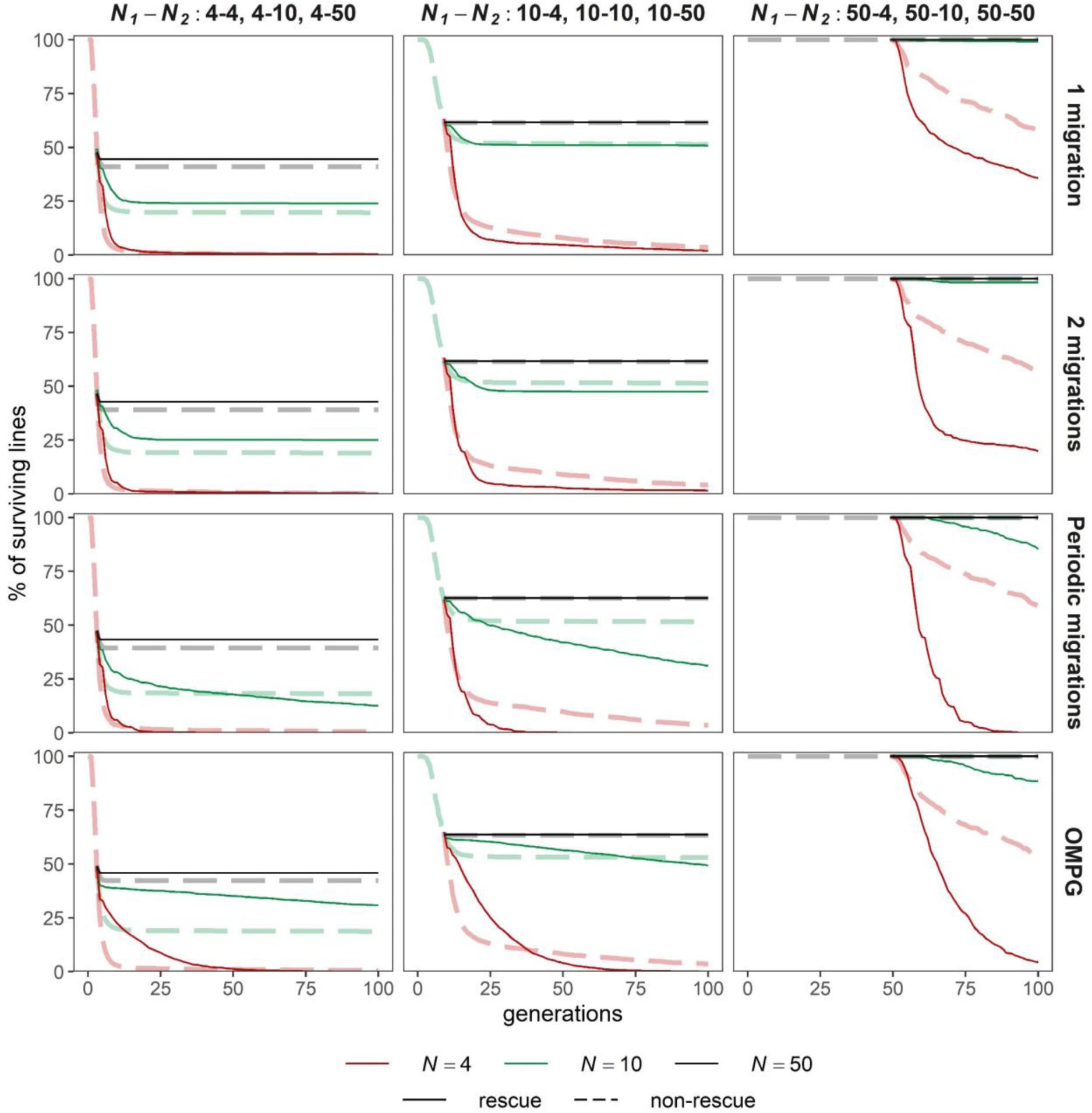
Percentage of surviving populations through generations. Different rows of panels give results under a different migration scenario: one unique migration event; two migrations with an interval of five generations; periodic migrations every five generations; “one migrant per generation” strategy. Different columns are for different *N*1-*N*2 demographic scenarios, coded as in Fig. Box. 1.2 panels. Light dashed lines give the percentage of surviving populations under no rescue program while solid lines give results under the rescue program. Number of migrants per event as explained in Box 1.

Under occasional migration, the rescue program increases the extinction risk in the cases where it induces a reduction of fitness compared to the non-rescue scenario (basically, when *N*_1_ > *N*_2_). Periodic migration and OMPG increase the extinction risk of small populations (*N* = 4 or 10) in the medium or long term even in cases where they cause an increase of fitness averaged over generations. The reason for this increased extinction risk is the genetic stochasticity introduced by migration events, as illustrated in Figure Box 1.3.

Figure Box 2.1 gives simulation results illustrating the effect of rescue programs on population survival. In these simulations, occasional migration after a history of severe census decline produced a small but relevant reduction of the accumulated extinction risk when coupled with an increase of the population size (scenarios 4-10 and 4-50). However, for very small populations where purging had been more efficient previously (scenarios 10-4 and 50-4), occasional migration caused more extinction risk than no migration.

Except for the largest population sizes (*N*_2_ = 50) where no further extinction occurred, periodic migration or OMPG increased extinction risk in the medium to long term, the reduction being more dramatic under periodic migration. Both strategies caused an increased extinction risk even in scenarios where they cause higher expected fitness. The reason is the stochastic nature of the introduced load, a phenomenon already noted in other simulation analyses (Robert et al. 2003). Under periodic migration this stochasticity adds to the periodic oscillations of fitness, due to accelerated inbreeding depression following the initial hybrid vigor after migration. The OMPG strategy removes the periodic component (Box 1.3), which makes more likely to reduce extinctions during some time. In both cases, each migration event introduced randomly sampled deleterious alleles leading to occasional abrupt fitness declines in individual populations, which can boost the risk of extinction. The fact that successive migration events favor extinction in cases where a single or two events improved survival, suggest that extinction occurs due to the fortuitous accumulation of successive random fitness declines in the same population, each fueled by a migration event where the sampled migrants harbored large load. Additional extinction results under other extinction criteria are shown in Figs. S9-S10, giving similar results to those reported above.

## Discussion

Our theoretical arguments and simulation results show that, considering a model of inbreeding load and genetic purging ascribed to partial recessive deleterious mutations, there are some specific situations where genetic rescue could be problematic. These situations are characterized by the use of migrants from non-purged donor populations that can introduce a substantial inbreeding load and genetic stochasticity into persistently small populations. The results suggest that additional caution needs to be introduced in the current genetic rescue paradigm (Ralls et al. 2018, 2020).

### Implications for conservation practice and some caveats of the simulation findings

In practical situations, the relevance of purging on the outcome of a rescue program depends on many circumstances that have not been addressed here, such as demographic and environmental stochasticity (particularly that affecting carrying capacity), adaptation to local conditions or the sex composition of migrants. Some of the factors not considered here can favor successful rescue. One main factor of hardly quantifiable consequences is the increased adaptive potential, which is favored by using large non-purged donor populations. Another possibly relevant factor is the introduction of favorable mutations occurred since the divergence between the donor and the recipient population, which are more likely to have accumulated in a large than in a small donor population (Ralls et al. 2020). This process of accumulation of new adaptive mutations is typically slow, so that it should not be relevant if the divergence is recent. On the contrary, if the divergence is more remote, the recipient population needs to have had large size during the majority of the divergence period in order to survive, so that it could have also accumulated different advantageous mutations, possibly implying some evolutionary divergence and risk of outbreeding depression (Edmands 2007). This is a subject that requires further investigation but, so far, there is little information available on the rate and nature of advantageous mutation for eukaryotes.

We have considered a model of inbreeding load *B* ascribed to the recessive deleterious component of many rare detrimental alleles that remain hidden in the heterozygous condition in a non-inbred population. This is the most parsimonious explanation of inbreeding depression for which estimates of mutation rates and distribution of effects have been widely found empirically, and has repeatedly been consistent with the analysis of laboratory evolutionary experiments (see, e.g., Caballero 2020, Chap. 8). We have not considered overdominance, which may also contribute to inbreeding depression, but whose relative contribution is speculative and, according to most evidence, probably minor (Charlesworth and Charlesworth 1999; Charlesworth and Willis 2009; Hedrick 2012; Thurman and Barrett 2016). In a reanalysis of some available estimates of genetic variance components for viability in *Drosophila melanogaster*, Charlesworth (2015) concluded that balancing selection should partly explain the excess of variance observed in some populations for viability with respect to mutation-selection balance (MSB) predictions. This conclusion, that could imply unrealistic low average viability (high segregating load) in large Drosophila populations (*i.e.*, high segregating load), is based on many crucial assumptions, including the supposition that *h* is fully determined by *s*, or that the additive variance ascribed to recessive deleterious alleles at large populations corresponds to the MSB expectation and can be predicted in terms of the average *h* value. However, the residual variability of *h* conditional on *s* can account for large amounts of inbreeding load and of variance, and recessive deleterious alleles can contribute substantially more additive variance in finite populations than expected at the MSB. No doubt that balancing selection due to antagonistic pleiotropy between fitness components can produce some excess in additive variance for viability (Fernández et al. 2005). However, an excess can also be expected in Drosophila due to pseudo-overdominance generated by linked deleterious mutations (see Waller 2021, for a recent study on the subject), because of the reduced length of the genome and its multiple inversions, as well as to genotype-environment interactions, as suggested by Mukai (1988) and Santos (1997). Contrary to the result of Charlesworth (2015), Sharp and Agrawal (2018) found no excess of variance for viability in laboratory Drosophila populations when compared to expected values computed using the decline of average viability in mutation accumulation lines. Although there was an excess of variance with respect to MSB predictions for fecundity and male mating success, it should be interpreted with caution due to i) natural selection possibly biasing the estimate of mutational mean decline in the mutation accumulation lines; b) possible differences between the magnitude of effects expressed during traits’ assay protocol and during population maintenance, as noted by the authors. In what follows, we concentrate on the consequences on rescue of the inbreeding load ascribed to deleterious alleles, and we left the consequences of overdominant loci to be explored in the future in cases where it might prove to be relevant.

In our simulations, migrant individuals were always males, but results would have been the same if migrants had been all females, because the distribution of the number of matings per individual, as well as that of the number of offspring contributed to the next generation, was the same for both sexes. However, in practical cases, using just female migrants can allow a better control of the amount and variability of inbreeding load introduced, while using males with a mating advantage can boost the short-term demographic rescue (Zajitschek et al. 2009) but also the spread of the immigrant’s inbreeding load, as in the case of the wolf’s population of Isle Royale (Hedrick et al. 2014, 2017 and 2019).

A consequence of all migrants having the same sex is that they always mate individuals of the endangered population. Therefore, a maximum hybrid vigor is expected in the first generation, but half of it would be expected to be lost in the absence of selection after one generation of panmixia. In the real world, the existence of maternal genetic effects on fitness may add a delayed component for hybrid vigor and inbreeding depression, obscuring the temporal fitness profile (Caballero 2020, p. 197). Therefore, the importance of the sex of migrants and of genetic maternal components on the dynamics of hybrid vigor and inbreeding load under repeated migration deserve being investigated.

Our finding of increased extinction risk for small populations under some scenarios of rescue programs seems to be in contradiction with the quite general view that, after excluding cases where outbreeding depression was to be expected, introduction of migrants always causes successful genetic rescue, usually assayed in terms of improved fitness or population sizes (Waller 2015; Frankham 2015, 2016; Ralls et al. 2020). However, evaluating genetic rescue effects is not simple (Robinson et al. 2020), and this view is grounded on rescue programs that had been tracked for just a few generations after immigration events (usually 1-3 generations, 5-6 on a few occasions) or on hybridization of populations (Chan et al. 2018), which is not the common situation in genetic rescue programs (Whiteley et al. 2015).

Our results are in disagreement in some respects with the simulation results obtained by Kyriazis et al. (2020), who found a substantial extinction rate for endangered populations with *N_e_* = 50 (or 25) which was always reduced under rescue programs. There are two main differences between the simulations by Kyriazis et al. (2020) and those presented here that could explain these disagreements. One is that they chose to simulate more realistic scenarios from an ecological point of view, in order to assess the joint consequences of genetic and nongenetic factors. We chose to illustrate the relevance of purging in simplified ecological conditions, an approach that allows a clearer understanding of the main genetic processes but lefts unexplored the relevance of their interaction with ecological factors. This could explain the larger extinction risks observed by Kyriazis et al. (2020). However, the different findings regarding the success of genetic rescue to prevent extinction is more likely to be due to the different mutational models used. Kyriazis et al. (2020), following a recent accepted trend, take the mutational model from estimates based on the evolutionary analysis of genomic data on site frequency spectra (Kim et al. 2017), which are very sensitive to the distribution of mild deleterious effects (*s* < 0.02 in homozygosis) but quite insensitive to the differences in deleterious effects above this threshold, which are conceptually pooled into a single “strongly deleterious” effect class. The problem is that, under these mutational models, the rate of mutations with *s* > 0.1 is tiny (Fig. 2A). However, there is evidence that a large fraction of the inbreeding depression that compromises the survival of endangered populations is due to large deleterious effects spread in the interval (0.1, 1], including lethals (Caballero and Keightley 1998; Bijlsma et al. 1999; García-Dorado et al. 2007; Fox et al. 2008; Charlesworth and Willis 2009; Hedrick et al. 2016; Domínguez-García et al. 2019). In addition, despite using a distribution of deleterious effects inferred assuming additivity, non-additive gene action is simulated. This is achieved by using a model equivalent to assigning *h* = 0.25 to any deleterious mutation with *s* < 0.02 and assigning *h* = 0 for *s* ≥ 0.02. This model is not consistent with inferences from classical experiments according to which deleterious mutations with *s* ≥ 0.02 are on the average only partially recessive (García-Dorado and Caballero 2000; Halligan and Keightley 2009; Ralls et al. 2020), as illustrated in Figure 2B. Under this mutational model, the inbreeding load concealed in each individual is ascribed to very many recessive deleterious alleles, each with a very small effect. Therefore, the coefficient of variation of the inbreeding load introduced by migrant individuals is very small, and the corresponding extinction risk ascribed to the genetic stochasticity is negligible.

**Figure 2.**
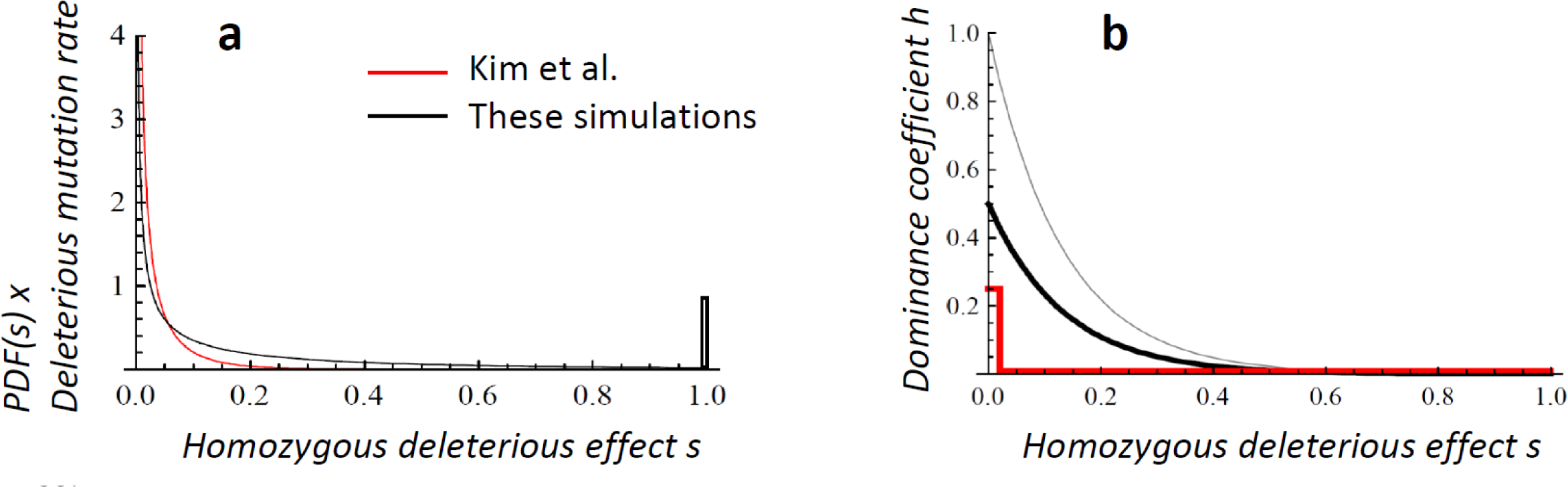
Mutational models. (a): Probability density function (PDF) of the homozygous deleterious mutational effects multiplied by the deleterious mutation rate. Red line: Model inferred from evolutionary genomic analysis by Kim et al. (2017) (best fit model for the 1000 genomes data: mutation rate per gamete and generation 0.314, homozygous effect *s* gamma distributed with shape parameter 0.186 and mean 0.0161, predicted equilibrium inbreeding load *B* = 3.07 for effective size 10^4^). Black line: Model used in our simulations (mutation rate per gamete and generation 0.2, *s* gamma distributed with shape parameter 0.33 and mean 0.2, predicted equilibrium inbreeding load *B* = 6.3 for effective size 10^4^); the lethal class generated in this model by assigning *s* = 1 to *s* values above 1 is represented in the [0.99-1] interval. (b): The black thick line gives the average inbreeding coefficient as a function of *s* assumed in our simulations, where *h* is uniformly distributed between 0 and the thin black line (extracted from García-Dorado and Caballero 2000 and García-Dorado 2003). The red line gives the *h* values used by Kyriazis et al. (2020) simulations, which are constant for each value of *s*.

In our simulations, we used a deleterious mutation rate and a joint distribution of *s* and *h* chosen to jointly account for: (a) the large rate of deleterious mutations unveiled by the evolutionary analysis of genomic data, with prevailing mild deleterious effect *s* < 0.01 that are likely to be roughly additive (Keightley and Eyre-Walker 2007; Boyko et al. 2008; Kim et al. 2017); (b) the results from classical mutation accumulation and fitness assay experiments, which are unlikely to detect small deleterious effects (say, *s* < 10^−3^) but imply a relevant rate of deleterious mutations with effects *s* > 0.1 that are severe from a conservation point of view (García-Dorado 1997; Caballero et al. 2002; Halligan and Keightley 2009; Thurman and Barrett 2016) and whose average degree of dominance is inversely related to their deleterious effect (García-Dorado and Caballero 2000; García-Dorado 2003); (c) the large inbreeding load concealed in large wild populations: under our deleterious mutation model, the expected haploid inbreeding load *B* for an equilibrium population with effective size *N* = 10^4^, computed by integrating the equation for the equilibrium inbreeding load (García-Dorado 2007) into the joint distribution for *s* and *h* was *B* = 6.23 (1.885 for lethal alleles), which is on the order of that found in several wild populations of mammals and birds (O’Grady et al. 2006; Hedrick and García-Dorado 2016); (d) The relatively large efficiency of genetic purging obtained from appropriate data under moderate bottlenecking (Bersabé and García-Dorado 2013; López-Cortegano et al. 2016). Fig. 2A illustrates that, under this model, the small rate of mutation with deleterious alleles that are severe in the conservation context (say, *s* > 0.1), is much larger than that assumed under the mutational model inferred from genomic evolutionary analysis. In our model, the dominance coefficient (Fig. 2B) is not completely determined by the homozygous deleterious effect although, according to empirical evidence, its expected value progressively decays with increasing *s*, so that most semilethal mutations are virtually recessive. Our knowledge on the joint distribution of mutational deleterious effects and dominance coefficients, including the spectra of severely deleterious effects above 0.1, is quite limited for species of conservation concern, so more information is needed in this respect. In any case, our results illustrate that it is necessary to be cautious, since the inbreeding load introduced by migrants from large non-purged populations can have an important sampling variance and the potential to compromise population survival depending on the demographic past and future of the endangered population.

### Considering the difference between reconnection and occasional or recurrent rescue

Our simulation results illustrate that continuous stable reconnection is safer than occasional or recurrent migration events. Furthermore, an important feature of reconnection in the wild is that population extirpation can in principle be reverted by recolonization, so that extinction should be referred to the metapopulation (Brown and Kodric-Brown 1977; Eriksson et al. 2014). Therefore, in this context, our simulation results for the evolution of fitness average under OMPG are more relevant than those for population survival which did not account for recolonization. Those fitness results suggest that continuous connection should improve the persistence of small populations. In practice, after restoring connectivity between the endangered population and a larger population or a metapopulation, the endangered population can enter an extirpation-recolonization dynamic that depends on many demographic and ecological factors (Franken and Hik 2004), as well as on the genetic stochasticity arising from the migration pattern. Then, an equilibrium is expected in the long term where the genetic identity of each population is scarcely affected by historical extirpation-recolonization events. In such situation, a stable but limited connectivity (*e.g.*, OMPG) can allow enough inbreeding from partial fragmentation to promote some purging, while preventing both the further progress of inbreeding depression and the metapopultion extinction.

Therefore, it is convenient to establish a conceptual distinction between genetic rescue programs based on occasional or even periodic migration, and programs aiming the continuous stable connection between the endangered population and a large healthy one (or a metapopulation). The former could be considered “*sensu stricto rescue*” programs, in the sense that they are intended to avoid the extinction of an isolated population whose stable reconnection is not feasible or whose differentiated genetic identity is worth to be preserved, although they involve some risk of swamping such identity. The second ones, say “*reconnection rescue*” programs, here represented by what might be considered its minimum migration rate (OMPG), may rather be aimed to preserve the endangered population as one of the valuable pieces integrating a metapopulation and the whole ecosystem, but one that does not show distinctive genetic features or adaptations to be preserved. This reconnection could be achieved either by actively moving individuals on a regular basis or by reconnecting landscapes, which has the advantage of setting an autonomous non-assisted mechanism. Besides allowing recolonization after extirpation, reconnecting landscapes will allow bidirectional flow, shifting the conservation aim from avoiding extinction of the endangered subpopulation to improving its long-term expected contribution to the metapopulation survival.

### The relevance of purging to rescue success under different conservation scenarios

The preceding sections show that, before introducing migrants from a large population into a critically endangered one, we should analyze the prospects that the increased reproductive potential expected from hybrid vigor immediately after migration and the habitat availability will allow the population to recover at least a moderate effective size in the near future, as in the case of Scandinavian wolves (Vilà et al. 2003). Furthermore, it is also convenient to gather information on the demographic history of the donor and the recipient population so that we can infer whether they underwent efficient purging in the past, as well as on the possibility that a stable reconnection can be achieved. To illustrate how our results can be considered to assist decisions regarding the introduction of migrants from a large non-purged population into an endangered one, below we present three different representative simplified scenarios.

1. In the first scenario, we consider the introduction of migrants to restore adaptive potential in an endangered population that has persisted for a long time with a moderate effective size (50 or above). In this case, the main purpose is not to reduce inbreeding depression or to ameliorate the load accumulated from continuous deleterious mutation, as both should be small due to past efficient purging (as well as to non-purging selection). Furthermore, if the population size does not decline further, purging is also expected to remove the load contributed by migrants, so that the genetic rescue program is not expected to be crucial by reducing inbreeding depression. In this situation, the main purpose of the rescue program is to restore adaptive potential and to prevent long term risk due to fixation of new deleterious mutation. Note, however, that inducing gene flow in populations that have survived for a very long time with moderate size and whose reproductive potential is large enough to allow population persistency may imply an unjustified risk, as has been appreciated in the case of Island foxes (Robinson et al. 2018). In particular, it could increase their inbreeding depression in case of future inbreeding. Special consideration deserves the case of endangered populations that have evolved differential adaptation or distinctive features, which could be swamped due to introgression or could lead to outbreeding depression (Hedrick and Fredrickson 2010; Frankham et al. 2011; Harris et al. 2019). However, there is some evidence that natural selection favoring locally adaptive alleles can efficiently prevent introgression of mis-adapted alleles during genetic rescue (Fitzpatrick et al. 2020).
2. In the second scenario, the rescue program is intended to restore both fitness and genetic diversity in a population that gets over a critical period of very small effective size (*N_e_* < 10) but has now recovered to some degree or is expected to do so quickly. During the past bottleneck, this population underwent high inbreeding, low purging and, therefore, an important reduction of fitness and of adaptive potential. The rescue program is expected to produce an immediate hybrid vigor and to provide wild alternatives to the deleterious alleles that might have been previously fixed in the endangered population. If the effective population size is moderately large at present (50 or above under our mutational model), or is expected to be so soon after receiving migrants, natural selection is expected to favor these introduced wild variants and to, slowly but efficiently, purge introduced recessive deleterious ones. And, of course, immigration will help to restore genetic diversity. Therefore, a rescue program integrated with measures that prompt early demographic restoration is expected to be helpful both in the short and the long term (Hufbauer et al. 2015). If the population recovery is less prosperous, one or two migratory events may be helpful to reduce the extinction risk before purging restores the population fitness, but periodic migration, advised in order to prevent the progressive loss of genetic diversity (Miller et al. 2020), could increase the extinction risk in the long term due to random accumulation of migration events introducing large amounts of inbreeding load. Then, a compromise should be achieved by favoring donor populations that are expected to be purged but that still contribute to enrich the genetic diversity of the recipient one.
3. Finally, the third scenario corresponds to rather extreme situations within the unfortunately paradigmatic case of an isolated population whose size has progressively declined over time, often beginning with a period where the decline was cryptic. With such demographic history, the ancestral inbreeding load should have been purged to a considerable extent. At present, the habitat has been reduced or degraded to a point that the effective size is dramatically small and is unlikely to grow in the near future. In such a situation, each migration event with individuals sampled from a large non-purged population produces some increase in mean fitness, but can be followed by accelerated inbreeding depression, increased stochasticity for fitness average, and increased extinction risk. The reason is that, due to the purging occurred while the size of the endangered population was moderate, the inbreeding load introduced by migrant individuals can be larger than the overall deleterious load of resident ones. Furthermore, the introduced load is boosted by the fitness advantage of the migrants and their vigorous crossed offspring (Saccheri and Brakefield 2002; Bijlsma et al. 2010), and it is inefficiently purged due to the small effective size. In these cases, periodically repeated rescue cycles allow the fortuitous concatenation of migration events introducing too high inbreeding load, worsening the survival prospects. Note that we are considering that, although the initial increase of fitness is expected to improve the intrinsic growth rate, the population size will continue to be small due to the limiting environmental factors and carrying capacity.

This scenario (iii) is vividly illustrated by the extinction of the Isle Royale wolves population, occurred after a sudden fitness increase caused by a single migrant from the continental population (Hedrick et al. 2019; Robinson et al. 2019). This scenario may seem of limited practical interest, as it refers to a very small effective population size. However, in wild populations, the effective size is usually much smaller than the actual number of breeding adults (Frankham 1995; Vucetich et al. 1997; Palstra and Ruzzante 2008). Furthermore, the effective size of many endangered populations has progressively declined down to values on the order of tens, as in the cases the Chatham Island black robin *Petroica traversi* (Ardern and Lambert 1997; Weiser et al. 2016) or the Fennoscandian arctic fox *Vulpes lagopus* (Angerbjörn et al. 2013; Norén et al. 2016), and many others. In some of these cases, population growth is limited by non-genetic factors, as habitat or prey limitations (Adams et al. 2011; Hedrick et al. 2014). Population growth can also be limited by inbreeding depression that could later be reverted to some extent due to genetic purging. Thus, this scenario, although extreme, can cover some practical conservation cases. Nevertheless, the specific risks in each situation are unknown due to the uncertainties surrounding the rate and effect distribution of deleterious mutation or the demographic history of populations, as well as to the many other genetic and non-genetic factors discussed above.

It is true that there is a pressure for taking action regarding critically endangered populations (Ralls et al. 2018), as their medium-term persistence is critically compromised if only due to demographic stochasticity. However, these results show that rescue interventions in persistently small populations may increase their long term extinction risk in some cases, calling for additional caution. In such cases, the rescue program should be coupled with reinforced habitat interventions in order to restore an effective population sizes large enough to allow efficient purging. Note that if, due to past purging under slow inbreeding, the endangered population shows no evidence of inbreeding depression, the rescue program is mainly intended to restore adaptive potential and might be postponed until a larger size is attained. Alternatively, one or several donor populations with a history of slow inbreeding, and therefore more purged, could be preferred, although the decision should weight the risk arisen from the introduced load with others derived from causes not included in these simulations, as the loss of adaptive potential. In any case, as far as the population singularity is not a main concern, the restoration of continuous connectivity should be preferred to recurrent migration events from time to time, due to fortuitous accumulation of random episodes introducing high load.

### Future advances and conclusions

The impact on conservation practice of the theoretical considerations and the simulation results discussed here depend on many factors that determine the genetic architecture of the inbreeding load, as the distribution of mutational effects and dominance patterns for deleterious mutations or the complexity of demographic histories, and all of them are worthwhile to be explored. Simulation approaches and experiments with model organisms can be very useful, both to advance in our understanding of this genetic features and to test the predictions generated in this analysis. It would also be helpful to understand how the genomic load assayed in terms of the burden of alleles annotated for different deleterious categories can inform on the fitness load and, in particular, on the inbreeding load measured in terms of concealed deleterious effects. Furthermore, there is a need for long term empirical observation of case studies, based on careful evaluation of the inbreeding load and the demographic and genetic flow history in both the donor and the recipient population, the evolution of fitness in the latter and the occurrence of extinction. These studies could benefit on the combined assay of fitness traits and genomic information.

The prospects of a rescue program depend on the demographic history of the endangered and donor populations but, in agreement with the small population paradigm, future population growth is essential to guarantee successful rescue and improve population survival. Our results illustrate that understanding all the consequences of conservation interventions is an arduous enterprise riddled with difficulties, and that the only safe strategy for *in situ* conservation and the one that should be prioritized and taken as a paradigm, is the recovery of large effective population size through the restoration of the habitat and of a healthy and continuous connectivity.

## Acknowledgements

We are grateful to the editor and to three anonymous reviewers by their helpful comments.

## DECLARATIONS

### Ethics approval

Not applicable

### Consent to participate

Not applicable

### Consent for publication

All authors have approved the manuscript for publication

### Availability of data and material

Not applicable

### Code availability

Codes will be available at GitHub address https://github.com/noeliaperezp/Genetic_Rescue

## SUPPLEMENTARY MATERIAL

### Methodology of simulations

We use computer simulations to explore the consequences of purging on genetic rescue programs considering different scenarios, always under mutation, natural selection and drift in a discrete generations model. First, we simulated a non-threatened population of *N* = 10^4^ dioecious diploid individuals with random mating for 10,000 generations in order to obtain a base population at the mutation-selection-drift equilibrium. Then, a smaller threatened population was derived and maintained until it was considerably inbred. The rescue program consisted in the introduction of a certain number of individuals from the large base population into the threatened one. Effective population size was assumed to equal the number of breeding adults (*N_e_* ≍ *N*).

### Simulation model and mutational parameters

Non-recurrent deleterious mutations occurred at rate *λ* = 0.2 per haploid genome and generation, with fitness effects being simulated through fecundity differences. For each locus, the fitness was 1, 1 – *sh*, 1 – *s* for the wild-type homozygote, the heterozygote and the homozygote for the mutant allele, respectively. The homozygous deleterious effect *s* was sampled from a gamma distribution with mean *s̅* = 0.2 and shape parameter *β* = 0.33, and the dominance coefficient *h* was obtained from a uniform distribution between 0 and e^(-*ks*)^, where *k* is a constant used to obtain the desired average value and ^ℎ̅^ = 0.283 (López-Cortegano et al. 2018). Thus, more deleterious alleles are expected to be more recessive (Caballero and Keightley 1994). Sampled *s* values larger than one were assigned a value *s* = 1 so that the mutational model generates a lethal class. The fitness of each individual was obtained as the product of genotypic fitnesses across loci. In order to produce each offspring, parental individuals were randomly chosen according to their fitness allowing for polygamous mating and free recombination. The haploid inbreeding load *B* for the base population (Morton et al. 1956), calculated as the sum of *s*(1 – 2*h*)*pq* for all selective loci, where *q* and *p* = 1 – *q* are the frequencies of the mutant and wild allele, respectively, was *B* = 6.23 (1.885 for lethal alleles), which is on the order of that found in several wild populations of mammals and birds (O’Grady et al. 2006; Hedrick and García-Dorado 2016). The number of segregating genomic sites with deleterious mutations was about 42,000 in the base population. In addition, 2,000 neutral sites, with reverse mutation allowed and the same mutation rate as sites under natural selection, were simulated to obtain estimates of neutral genetic diversity.

### Threatened populations and rescue program

Different scenarios were simulated as shown in Fig. Box 1.1 of the main text. In a first phase, threatened populations with different sizes (*N*_1_ = 4, 10 or 50) were maintained under the same conditions as the base population except in that offspring were randomly assigned to male or female sex with equal probability. A second phase started at generation *t* = *N*_1_ (e.g., at generation 50 for populations with *N*_1_ = 50, etc.) so that the expected average inbreeding coefficient was *F* ≍ 0.4. At this point, four alternative scenarios were simulated for each threatened population (second phase, with population size *N*_2_; Table S1), where the population was maintained with the same constant size (*N*_2_ = *N*_1_) or with a different constant size (*N*_2_ ≠ *N*_1_), and entered or not a genetic rescue program. Regarding the population size in these two phases, these scenarios will be denoted by the corresponding numbers (*e.g*., 50-10 stands for threatened populations with population size 50 during phase 1 and 10 during phase 2). In order to enforce mating between native and migrant individuals, these were assumed to be males. Migrants did not replace the individuals of the line (*i.e*., the number of individuals after a migration event was the size of the line plus the number of migrants). The whole scheme was simulated in a single round, so that each set of four scenarios shared the same original threatened population. Depending on the case, between 500 and 2,500 replicated rounds were simulated. The number of individuals introduced during each migration event was five for lines with *N*_2_ = 50 and one for lines with *N*_2_ < 50. Regarding the number of migration events, we considered four strategies: i) a single event; ii) two events with an interval of five generations; iii) periodic migration every five generations; iv) the “one migrant per generation” (OMPG) strategy, that is widely recommended to retain connectivity in metapopulation management (Mills and Allendorf 1996). All the sizes considered for the threatened populations (4, 10 and 50) correspond to the IUCN Red List category of Critically Endangered or Endangered according to Criterion D (IUCN 2012). Extinction of a line occurred when the average fitness (*w*) of males and/or females was less than 0.05 and/or when there were only breeding males or only breeding females (counted after migration when appropriate). Alternative criteria assumed extinction when *w* = 0 or *w* < 0.1, or no extinction at all.

**Table S1.**
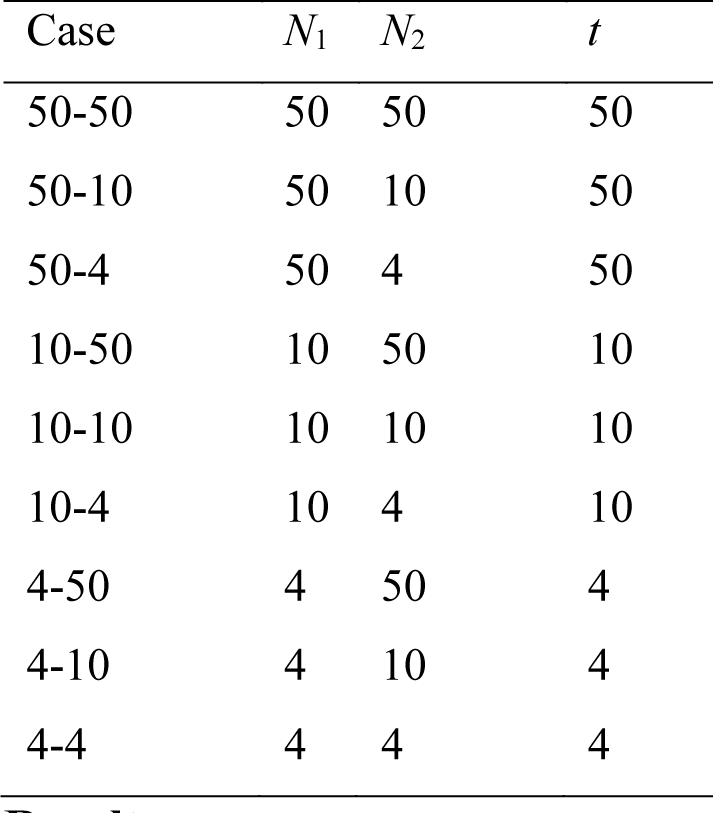
Simulation scenarios. *N*_1_: population size of the threatened population during the first phase. *N*_2_: population size of different scenarios derived from the initial threatened population (phase 2). *t*: generation at which *N*_1_ is modified to *N*_2_ and some populations enter a genetic rescue program.

| Case | $N_1$ | $N_2$ | $t$ |
| --- | --- | --- | --- |
| 50-50 | 50 | 50 | 50 |
| 50-10 | 50 | 10 | 50 |
| 50-4 | 50 | 4 | 50 |
| 10-50 | 10 | 50 | 10 |
| 10-10 | 10 | 10 | 10 |
| 10-4 | 10 | 4 | 10 |
| 4-50 | 4 | 50 | 4 |
| 4-10 | 4 | 10 | 4 |
| 4-4 | 4 | 4 | 4 |

Each generation we computed average fitness (*w*), genetic diversity (*H*) for the neutral loci, and overall inbreeding load (*B*), always excluding migrants.

## Results

### Evolution of threatened populations without genetic rescue

Figures S1-S4 give the evolution of different genetic parameters averaged over replicates under different scenarios assuming no extinction (results for a subset of scenarios are given in Fig. Box 1.2). Figures S5-S8 give analogous results obtained for the set of surviving lines, which are qualitatively similar to Figs. S1-S4 except in that fitness averages are a little higher when extinction is high, and in that the sampling error of the over-replicates averages for the different parameters increases as the number of surviving lines drops.

Results on the percent of surviving lines, analogous to those presented in the main text but assuming different extinction criteria are shown in figures S9 and S10. Results are similar under the different extinction criteria.

---

**Figure S1.**
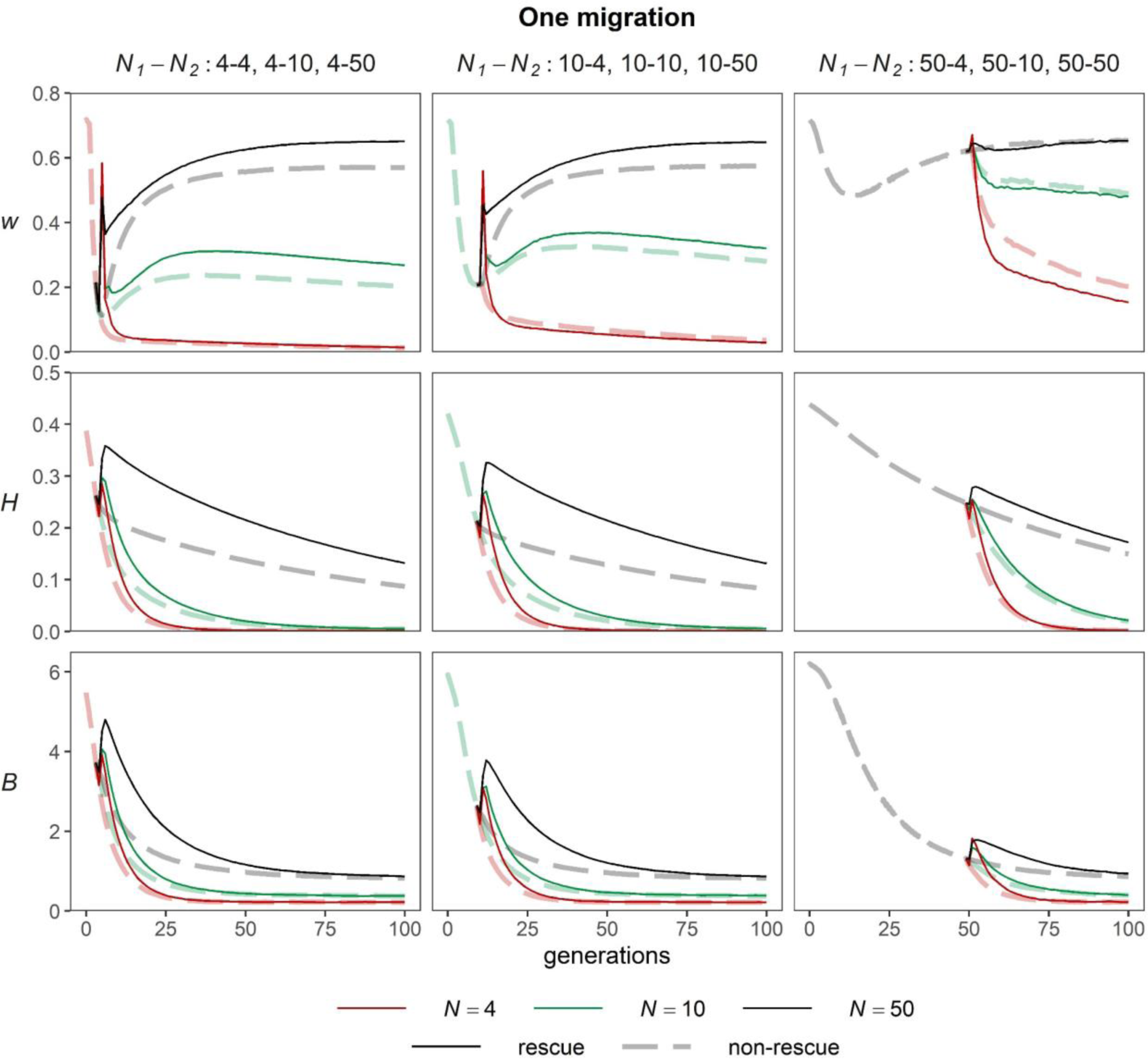
Evolution of fitness (*w*), genetic diversity (*H*) and inbreeding load (*B*) of threatened populations entering a genetic rescue program (one unique migration of 5 individuals in lines *N*_2_ = 50, and 1 individual otherwise; solid lines) and of control threatened populations (dashed lines). No extinction allowed. To avoid extinction due to all breeding individuals being homozygous for lethal alleles, we assigned *s* = 0.99 whenever the *s* value sampled from the gamma distribution was larger than 0.99 (the standard procedure in the cases with extinction allowed was assigning *s* = 1 when the sampled value was larger than 1).

**Figure S2.**
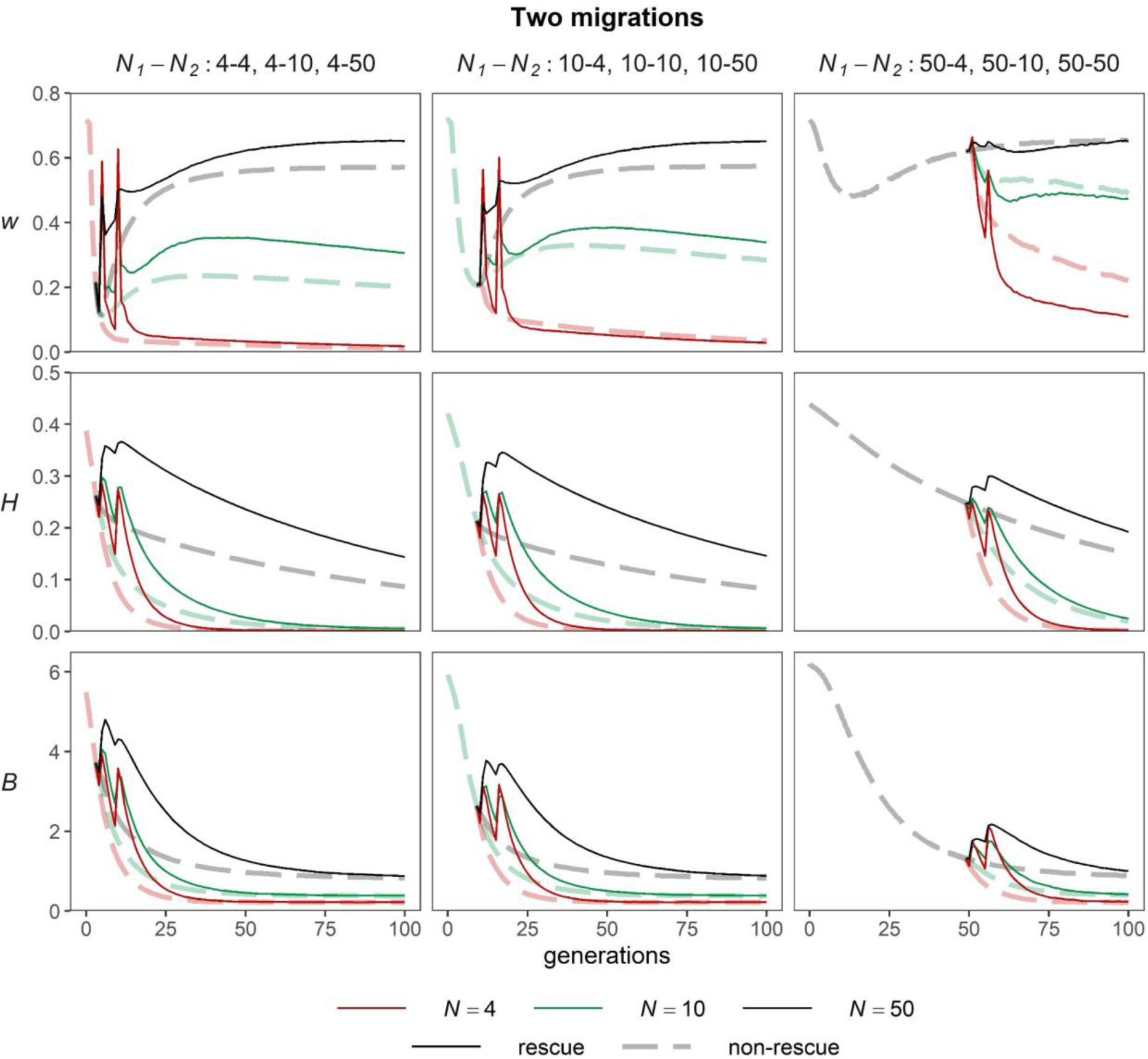
Evolution of fitness (*w*), genetic diversity (*H*) and inbreeding load (*B*) of threatened populations entering a genetic rescue program (two migrations of 5 individuals in lines *N*_2_ = 50 with an interval of five generations, and 1 individual otherwise; solid lines) and of control threatened populations (dashed lines) without extinction (as in Figure S1).

**Figure S3.**
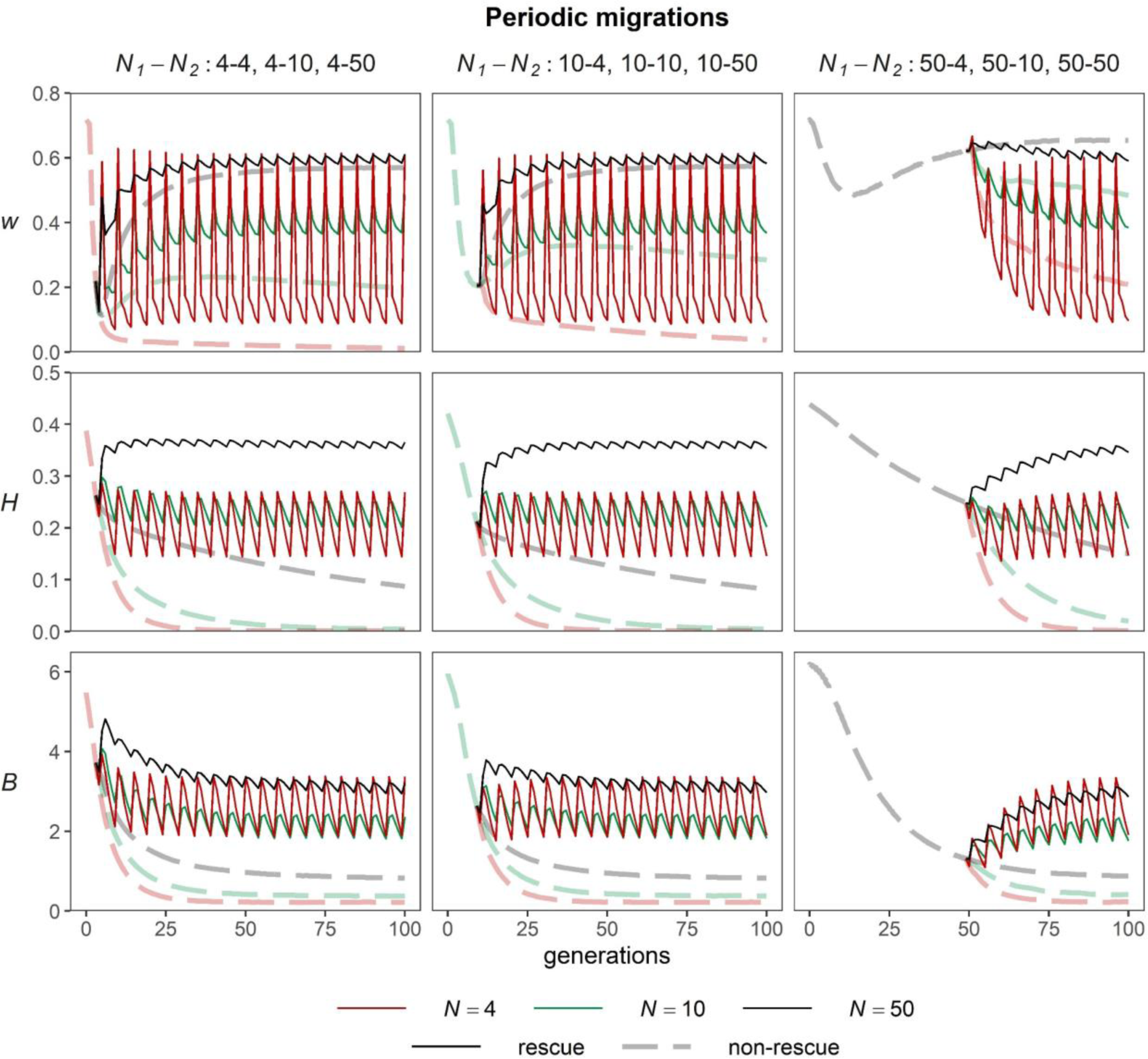
Evolution of fitness (*w*), genetic diversity (*H*) and inbreeding load (*B*) of threatened populations entering a genetic rescue program (periodic migrations every five generations of 5 individuals in lines *N*_2_ = 50, and 1 individual otherwise; solid lines) and of control threatened populations (dashed lines) without extinction (as in Figure S1).

**Figure S4.**
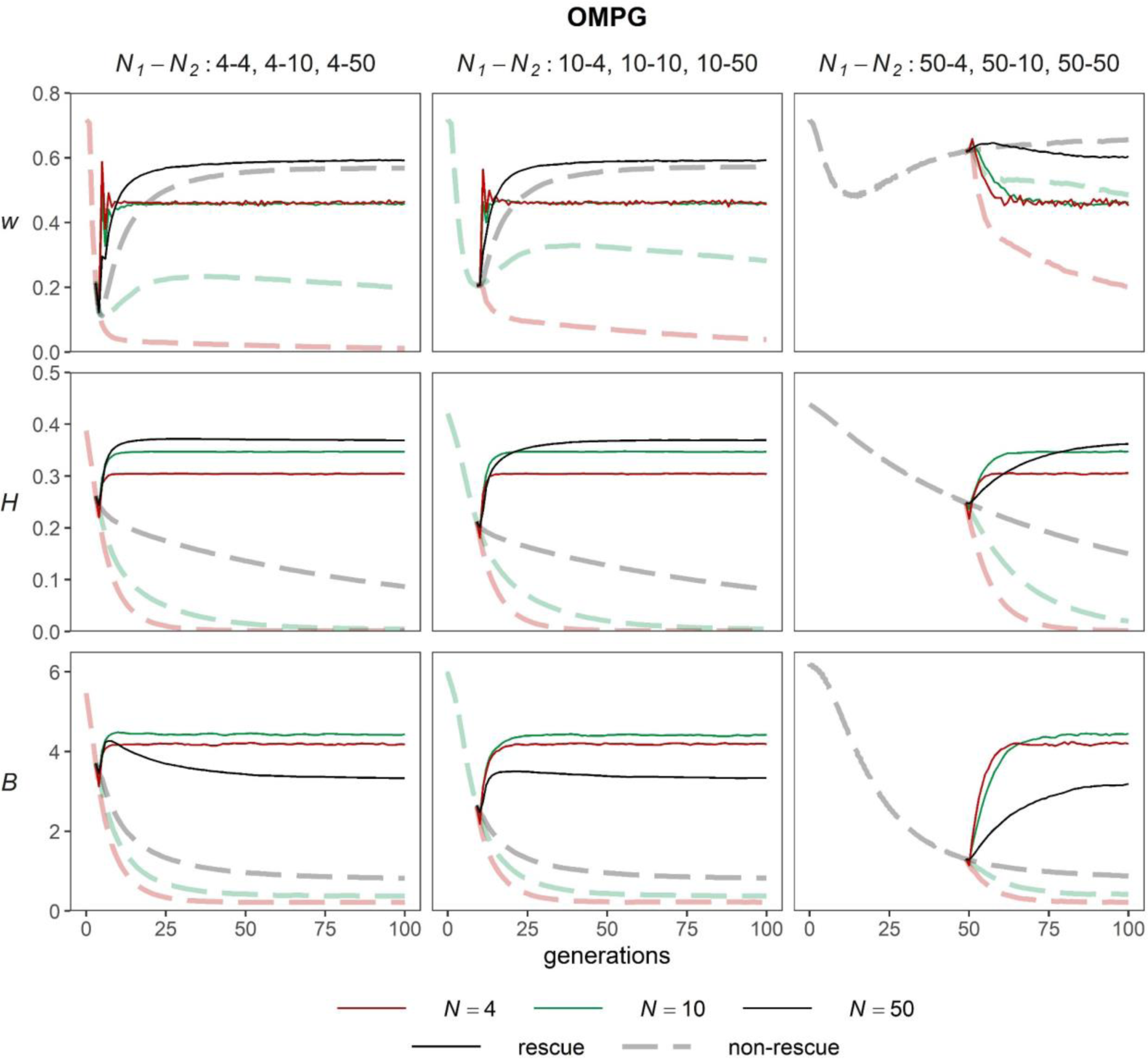
Evolution of fitness (*w*), genetic diversity (*H*) and inbreeding load (*B*) of threatened populations entering a genetic rescue program (“one migrant per generation” strategy; solid lines) and of control threatened populations (dashed lines) without extinction (as in Figure S1).

**Figure S5.**
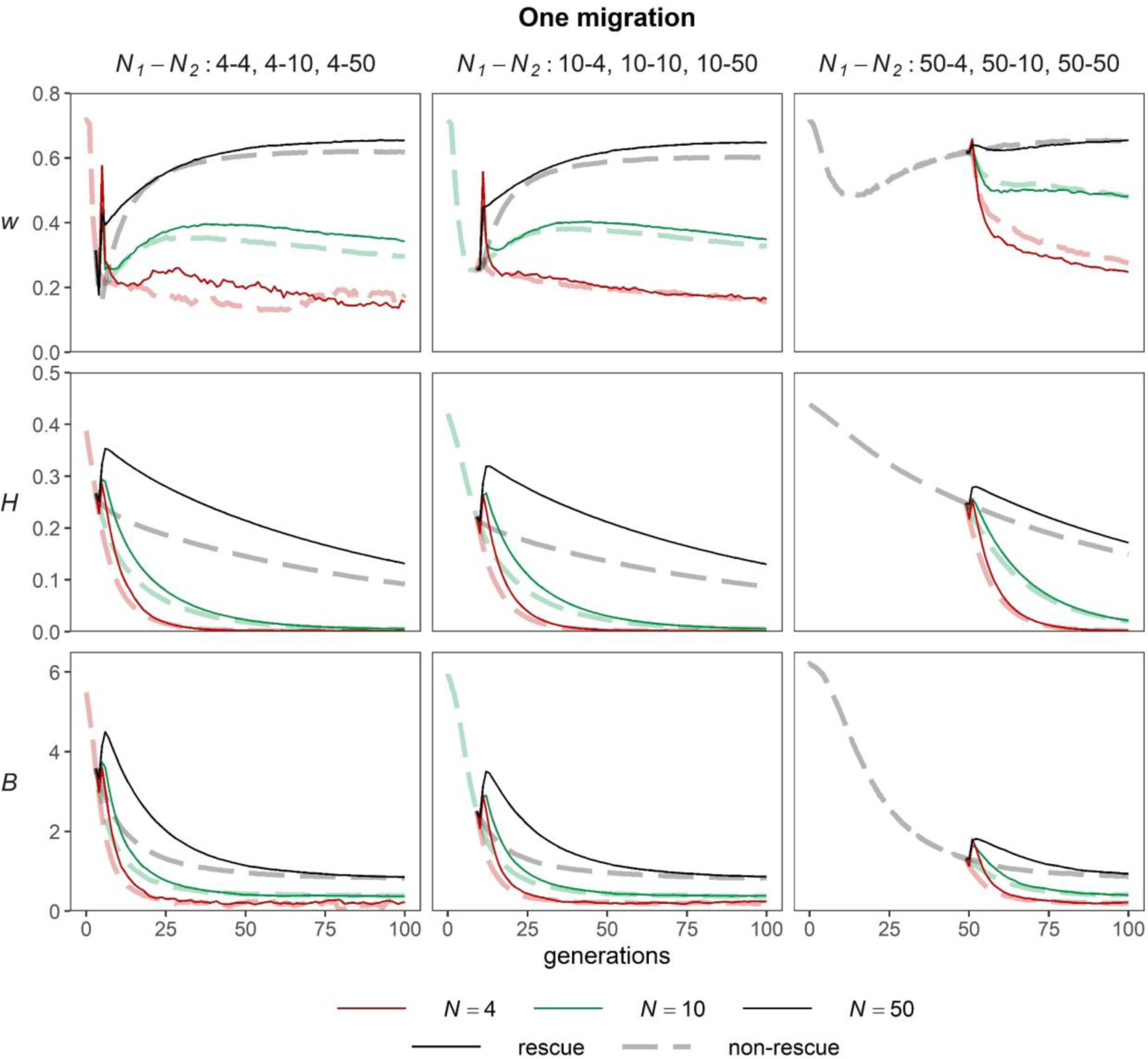
Evolution of fitness (*w*), genetic diversity (*H*) and inbreeding load (*B*) of threatened populations entering a genetic rescue program (one unique migration of 5 individuals in lines *N*_2_ = 50, and 1 individual otherwise; solid lines) and of control threatened populations (dashed lines). Extinction of a line occurred when the average fitness (*w*) of males and/or females was less than 0.05 and/or when there were only breeding males or only breeding females.

**Figure S6.**
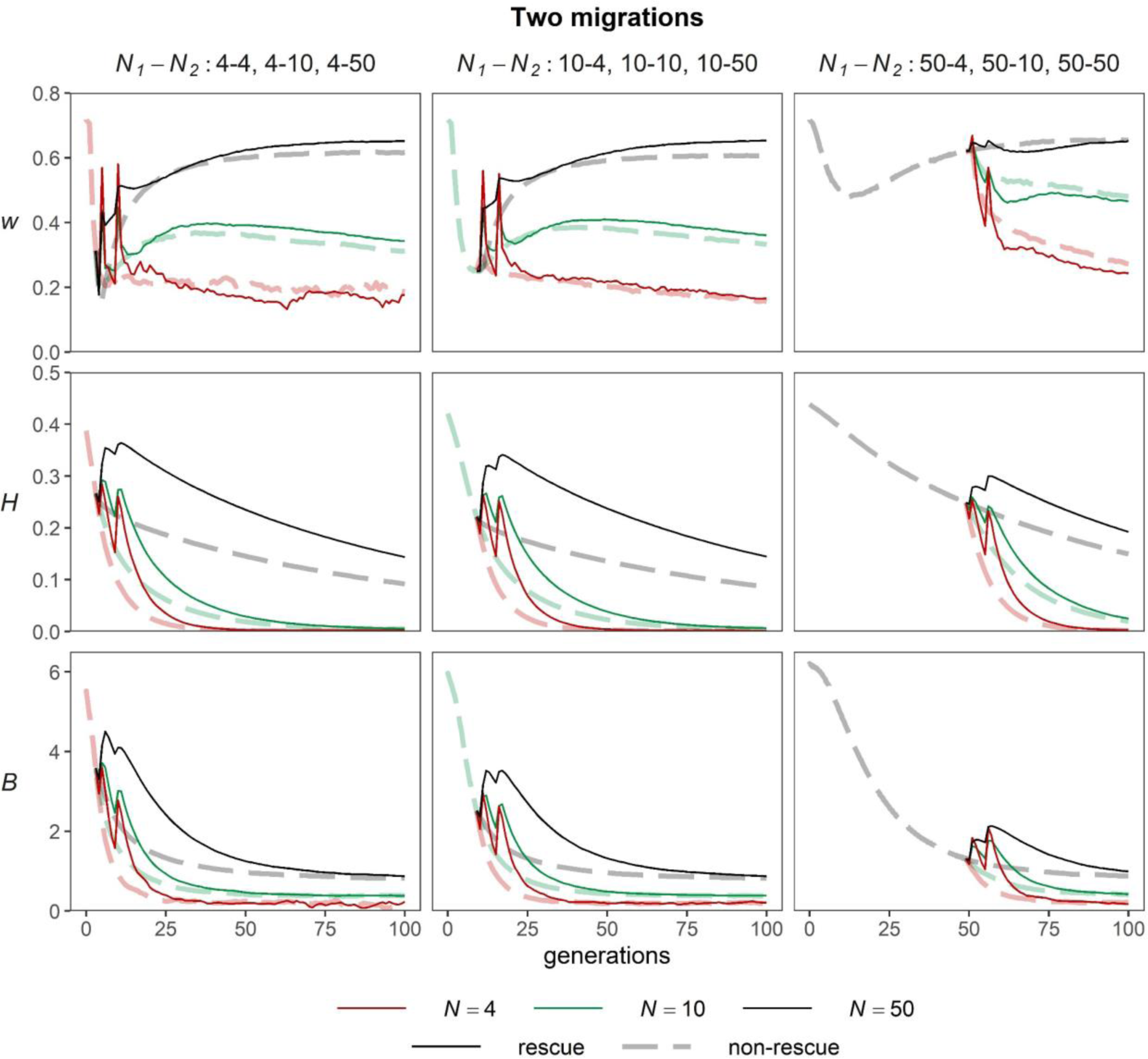
Evolution of fitness (*w*), genetic diversity (*H*) and inbreeding load (*B*) of threatened populations entering a genetic rescue program (two migrations with an interval of five generations of 5 individuals in lines *N*_2_ = 50, and 1 individual otherwise; solid lines) and of control threatened populations (dashed lines). Extinction of a line occurred when the average fitness (*w*) of males and/or females was less than 0.05 and/or when there were only breeding males or only breeding females.

**Figure S7.**
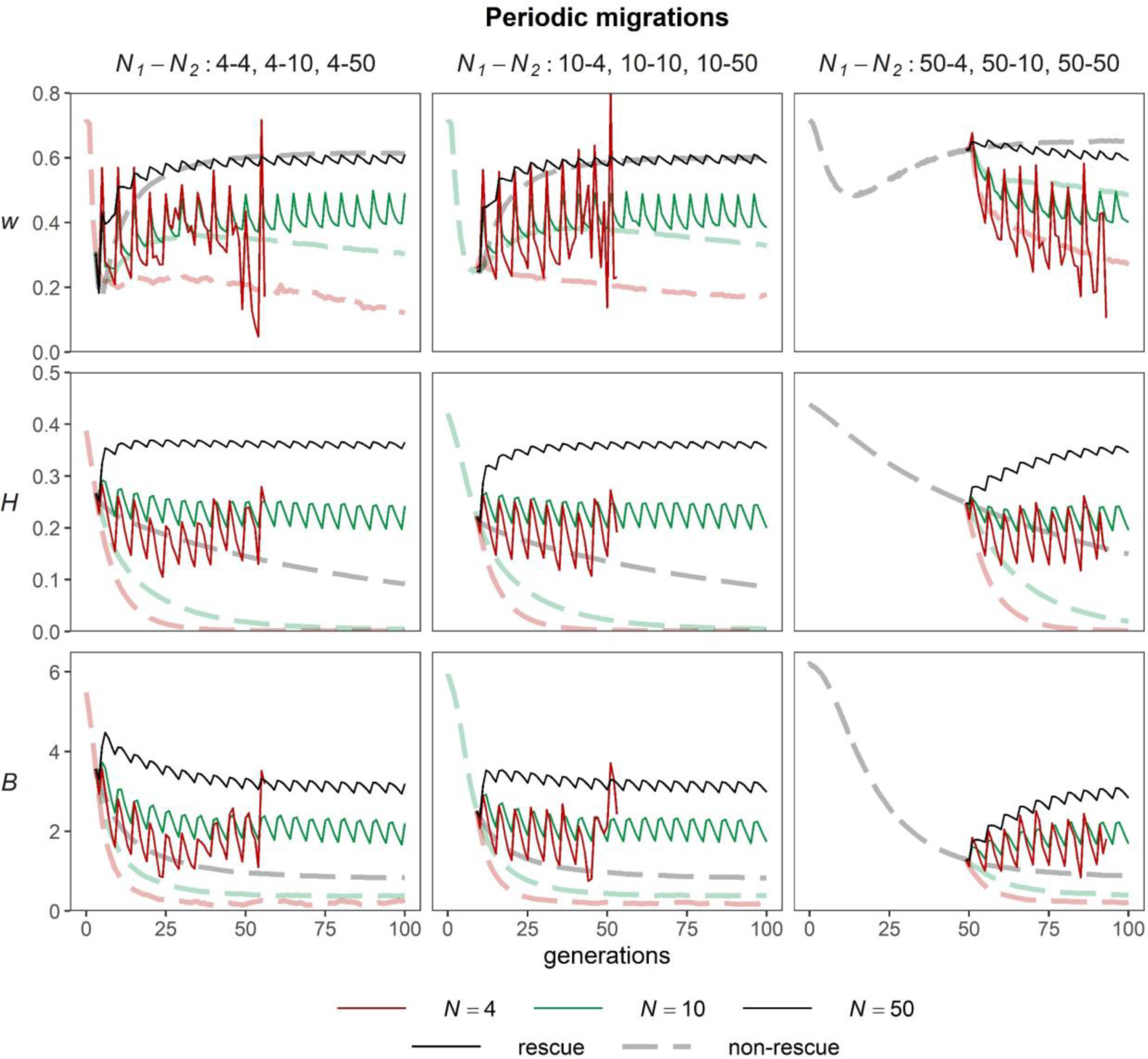
Evolution of fitness (*w*), genetic diversity (*H*) and inbreeding load (*B*) of threatened populations entering a genetic rescue program (periodic migrations every five generations of 5 individuals in lines *N*_2_ = 50, and 1 individual otherwise; solid lines) and of control threatened populations (dashed lines). Extinction of a line occurred when the average fitness (*w*) of males and/or females was less than 0.05 and/or when there were only breeding males or only breeding females.

**Figure S8.**
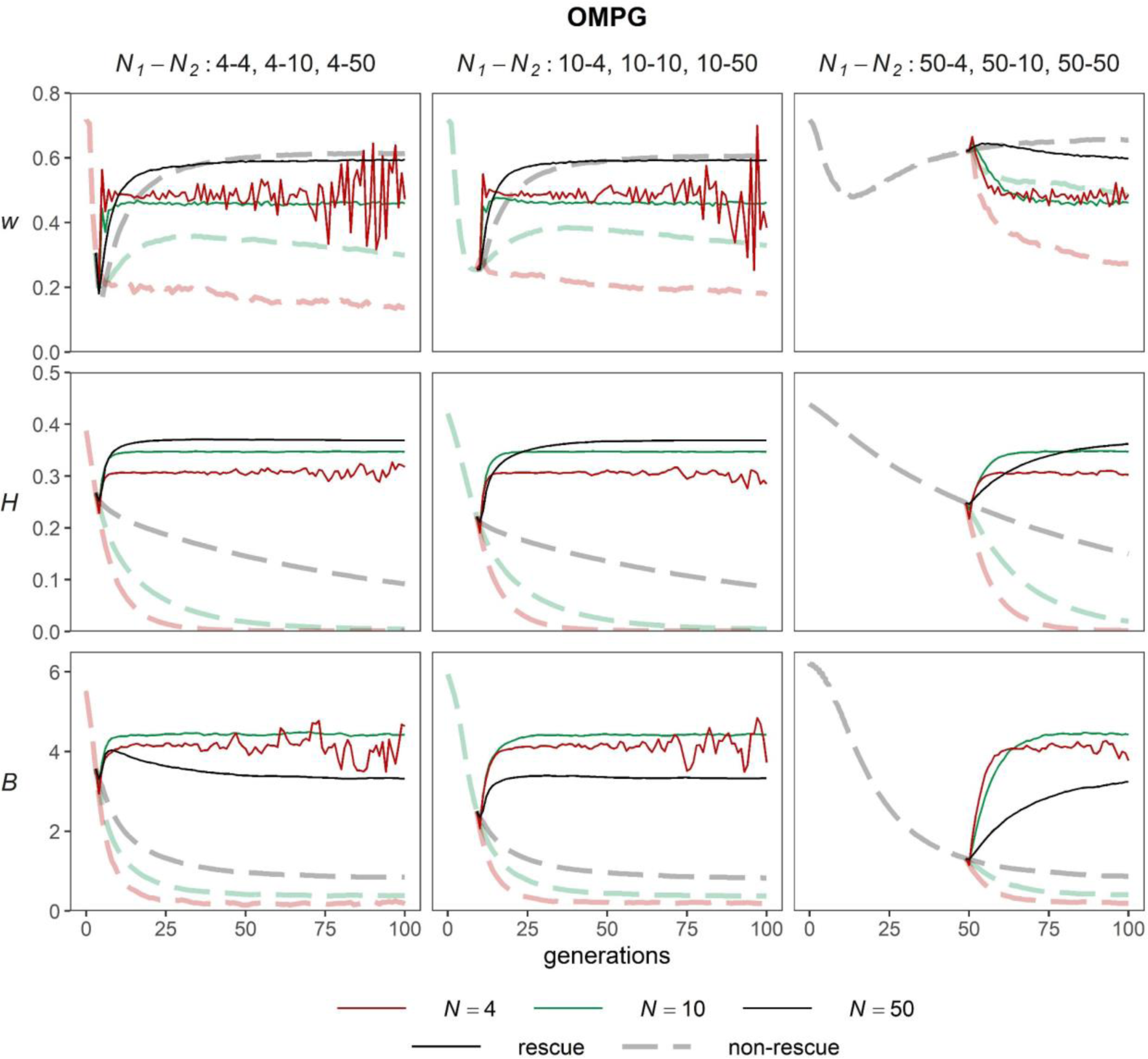
Evolution of fitness (*w*), genetic diversity (*H*) and inbreeding load (*B*) of threatened populations entering a genetic rescue program (“one migrant per generation” strategy; solid lines) and of control threatened populations (dashed lines). Extinction of a line occurred when the average fitness (*w*) of males and/or females was less than 0.05 and/or when there were only breeding males or only breeding females.

**Figure S9.**
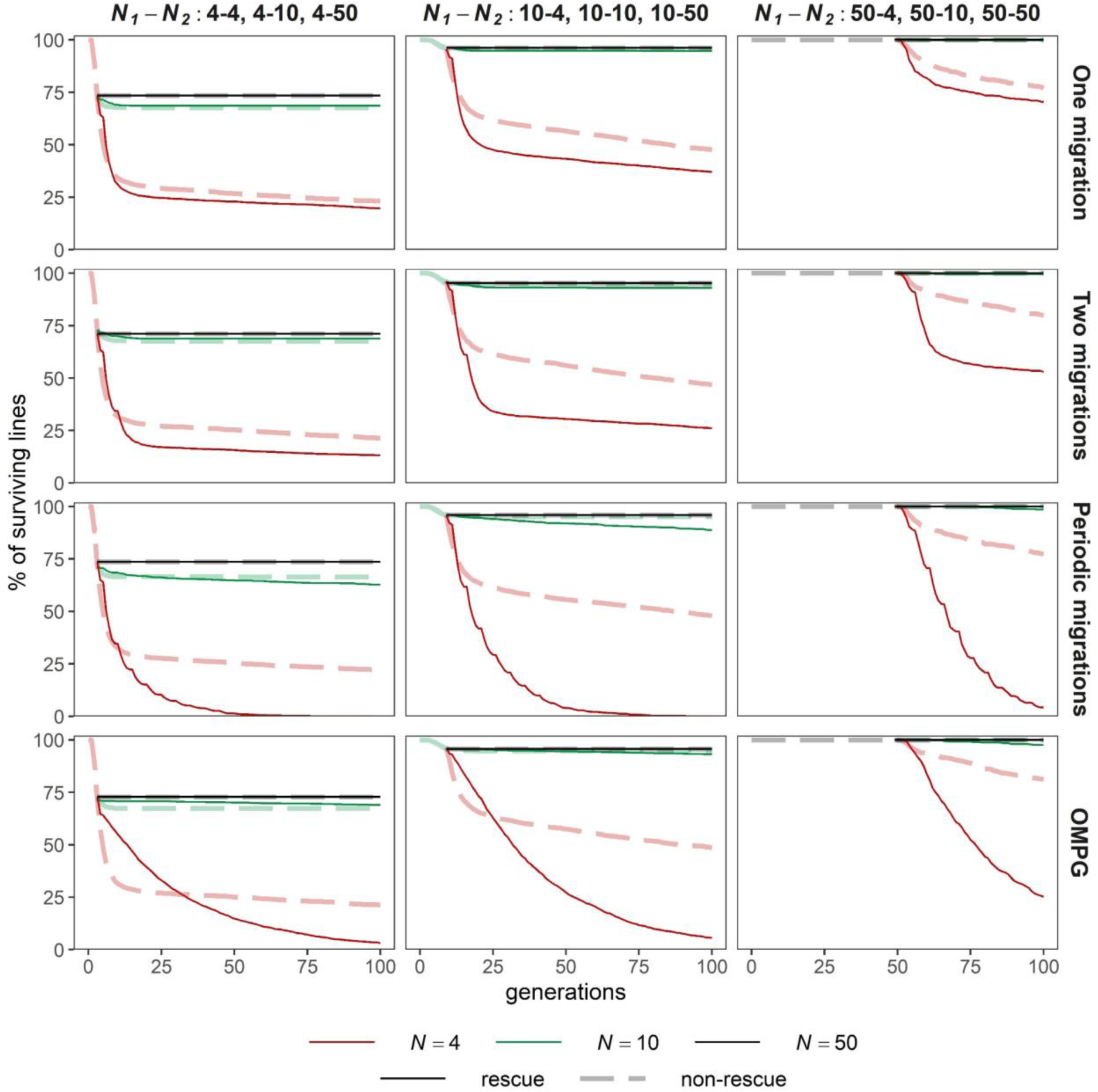
Percent of surviving threatened populations under a genetic rescue program (one unique migration; two migrations with an interval of five generations; periodic migrations every five generations; “one migrant per generation” strategy; solid lines) and of surviving control threatened populations (dashed lines). Extinction due only to homozygosis for lethal alleles (*i.e.*, extinction occurs when fitness is 0 for all the males or/and all the females in the line) or to all breeding individuals being of the same sex.

**Figure S10.**
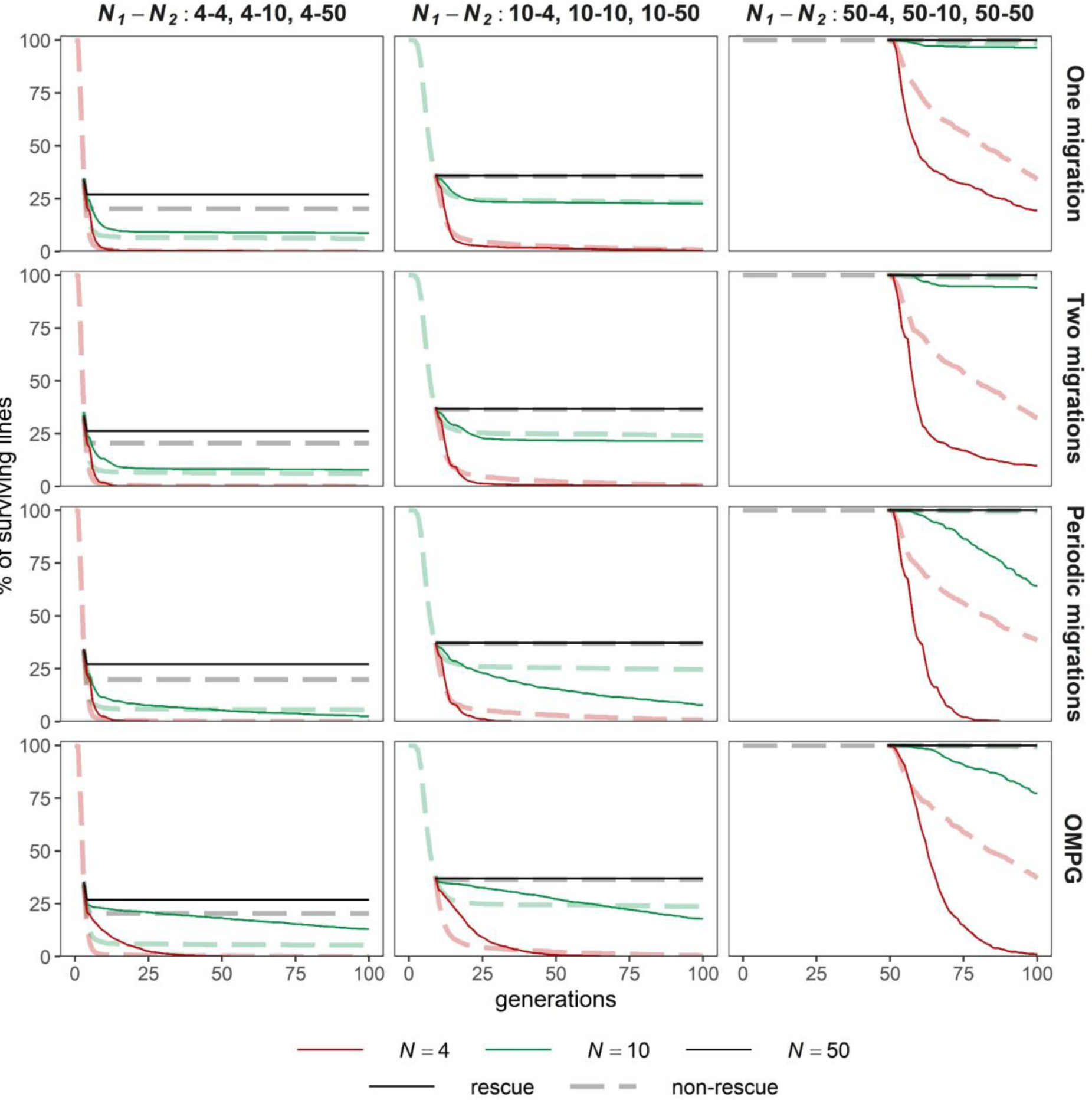
Legend as in Figure S9, but extinction occurs when the average fitness (*w*) of males and/or females is less than 0.1.

